# Intraflagellar transport of tubulin maintains steady-state axoneme integrity in *C. elegans* cilia

**DOI:** 10.64898/2026.04.14.718528

**Authors:** Elizaveta Loseva, Aniruddha Mitra, Daniël Groskamp, Erwin J.G. Peterman

**Author notes:** Corresponding author: Erwin J.G. Peterman.

## Abstract

The integrity of the axoneme – the microtubule (MT)-based core of the cilium – and intraflagellar transport (IFT) are interdependent. The mechanisms determining axoneme structure and dynamics have remained largely unknown, especially in primary cilia with a more variable architecture and longer MT singlet parts. Using fluorescence imaging in the phasmid neurons of *C. elegans*, we here demonstrate that β-tubulin isotype TBB-4 diffuses through the dendrite and employs a combination of anterograde IFT and diffusion to reach the sites of incorporation in the steady-state axoneme. Disrupting tubulin’s ability to bind to the IFT significantly reduces its share in the axoneme. We suggest that, in phasmid cilia, a constant supply of tubulin by IFT is required for steady-state length maintenance, in order to elevate soluble tubulin concentration near the axonemal tips and to promote MT stability.

## Introduction

Cilia are elongated organelles protruding from the cell surface of many eukaryotic cells. Motile cilia, or flagella, drive cellular motion or propel fluids surrounding the cell. Primary, or sensory cilia, appear to have originated from motile cilia but have lost their ability to beat and have evolved into specialised cellular antennae capable of receiving and transducing environmental signals [1]. Owing to their common evolutionary origin, these two functionally distinct types of cilia share largely similar inner architectures and underlying molecular mechanisms [1, 2].

The core structure of the cilium, the axoneme, consists of 9 radially oriented microtubule (MT) doublets originating from a basal body, a modified centriole. Motile cilia also possess a central pair of singlet microtubules (commonly designated as the “9+2” architecture), as well as radial spokes, nexin links and axonemal dynein arms, connecting the MTs and providing the beating ability [3-5]. Axonemes of primary cilia have initially been described as analogous to the ones of motile cilia, but missing the contractility-associated elements and the central pair of microtubules (and are consequently designated “9+0”). However, later studies have revealed a high degree of variability in axonemal structures: in different cell types, primary cilia can possess a variable number of additional singlet MTs, MT doublets can transition into relatively long singlets, twist, splay and so on [6-8], which are designated as “9v” (v for variable) [9]. Proteins associated with axonemal MTs also differ between motile and primary cilia [6, 10, 11]. Unlike dynamic cytoplasmic MTs, axonemes in both primary and motile cilia are typically stable, with only plus-tips undergoing tubulin exchange [12, 13]. Cells can resorb or shed their cilia, e.g. before mitosis or in response to external stimuli, and then regrow cilia to their initial lengths [14, 15].

Ciliary integrity and function are maintained by intraflagellar transport (IFT), a specialised active transport system in which large polymers of protein complexes, called IFT trains, move back and forth along the axoneme, powered by kinesin-2 motors in the anterograde direction (towards the ciliary tip) and dynein-2 in the retrograde direction (towards the base) [16]. Cargo molecules, such as receptors, signalling cascade actors, or cytoskeleton components, bind to the IFT trains, either directly or via adapter complexes, to reach their destination point in the cilia [17-22].

Tubulin, the main component of the axoneme and by far the most abundant protein in the cilium, also partially depends on IFT for its localisation. One experimentally confirmed bipartite tubulin binding site is formed by the N-terminal domains of two IFT-B core proteins, IFT81 and IFT74 [23]. IFT81N forms a calponin homology (CH) domain capable of binding the globular parts of an αβ-tubulin dimer, while unstructured, highly basic IFT74N domain forms electrostatic interactions with the acidic C-terminal tail of β-tubulin. Both sites contribute to tubulin binding, and while their affinities for tubulin could not be measured independently, IFT74N is supposedly the major contributor, based on the severity of the effects caused by the respective mutations. A recent cryo-ET-based reconstruction of anterograde IFT trains has revealed that such canonical tubulin binding with IFT81CH appears to sterically interfere with neighbouring IFT-B complexes [24]. One hypothesis explaining this apparent contradiction could be that tubulin binds to the IFT81/IFT74 complex at the ciliary base, but that during the anterograde IFT train assembly, a structural change causes IFT81N to dissociate from tubulin, leaving IFT74N as the only binding site [25]. In line with that, IFT74N phosphorylation, weakening the electrostatic interactions with β-tubulin C-terminal tail, has been suggested to trigger tubulin unloading in the cilia of *C. elegans* [26].

In the biflagellate green algae *C. reinhardtii*, tubulin transport by IFT has been shown to be upregulated in growing cilia, while it becomes almost exclusively diffusive in steady-state cilia, despite IFT train frequency being unaffected [27]. The mode of tubulin transport is supposedly regulated by the concentration of soluble tubulin in the cell body at different cell stages, although additional regulating factors might play a role as well. During the fast phase of flagellar growth, one tubulin-binding site per IFT-B complex can provide only ~25% of the required tubulin [28]. Several other IFT proteins with CH domains have been suggested as additional binding sites [28], one of which was shown to bind tubulin *in vitro* [29]. However, disrupting both interactions within the IFT81/IFT74 binding site has been shown to halt tubulin IFT by 99% [30], so there is no experimental evidence for substantial tubulin binding elsewhere on the IFT train. Remarkably, while tubulin-binding-associated mutations in the IFT81/IFT74 module result in the formation of very short, truncated axonemes [31], disrupting this IFT-tubulin interaction from the tubulin side results only in a slight, ~17% reduction of its share in the axoneme [30], leaving the possibility that mutating both IFT81N and IFT74N might affect some aspects of ciliary maintenance other than tubulin transport. The notion that IFT has insufficient capacity for tubulin, together with the only mild effects of disrupting tubulin transport by IFT, has been interpreted to suggest that most tubulin in the cilia is supplied by diffusion, which was also confirmed by high-speed imaging [30]. Collectively, these data support a model in which the principal role of IFT in axoneme maintenance is to concentrate tubulin at the MT tips, facilitating axonemal growth.

The insights discussed above have been obtained for the motile cilia of *C. reinhardtii*. Analogous studies in primary cilia have remained rather elusive. Studies utilising FRAP, *in silico* modelling and high-speed super-resolution microscopy have suggested that tubulin undergoes bidirectional active transport, as well as diffuses in the axonemal lumen of primary cilia [12, 32]. However, direct observation of tubulin trajectories has not been achieved. In human cell lines, removal of IFT81N causes a complete loss of cilia, a phenotype similar to that of the IFT81-null mutation [23]. This effect is more severe than in motile cilia, where disrupting one binding site on the IFT81/IFT74 module only moderately affects tubulin transport frequency and flagellar regeneration rate [31]. This might indicate that tubulin IFT is more important in primary cilia than in motile one. It is however important to note that in the human cells, the mutation in IFT81 involved the removal of a complete protein domain, rather than only a few amino acids, which might also be the cause of the more severe phenotype observed.

Here we present new insights from ensemble and single-molecule dynamics of β-tubulin isotype TBB-4 in the phasmid chemosensory cilia of *C. elegans*. By exploiting local photobleaching and high-speed fluorescence imaging, we observed both active anterograde transport and diffusion of TBB-4 and found that, unlike in the motile cilia, tubulin transport by IFT is also active during steady-state in primary cilia. Further, we reveal that transport of tubulin as cargo of IFT trains is key for its localisation near the axonemal tip and hypothesise that IFT-driven transport is key for maintaining the stability of the dynamic tips of the axoneme. This might reflect fundamental differences in axoneme stability between motile and primary cilia.

## Results

### TBB-4::eGFP is present all over phasmid neurons, except for the ciliary base, where it is depleted

To explore the dynamics of axonemal tubulin, we performed fluorescence microscopy on *C. elegans* strains expressing eGFP-labelled β-tubulin isotype TBB-4, using a Mos-1-mediated single-copy insertion strain (MosSCI) in which the original copy of the tbb-4 gene was still present [33]. In fluorescence images of the worm’s tails, we observed that TBB-4::eGFP was mostly expressed in five ciliated neurons: PHA (L, R), PHB (L, R) and PQR. TBB-4::eGFP is visible throughout these cells and appears enriched in the cilia (Figure 1A). Comparing the (integrated) fluorescence intensities in such images to that of single eGFP molecules (Supplementary Figure 1) allowed us to estimate that each PHA/PHB cilia pair (L or R) contains (8.0 ± 1.7) × 10^3^ TBB-4::eGFP proteins (average ± error estimated using bootstrapping (see methods); N = 11 cilia). The total number of β-tubulins incorporated in the axonemes of a pair of cilia is expected to be about 346 500 (assuming that each axoneme consists of 9 doublet MTs of 5 µm length and 9 singlet MTs of 3 µm, ignoring free tubulin in the cilium). From the fluorescence intensities, we can thus estimate that 2.3 ± 0.52% of the total β-tubulin in the axonemes is TBB-4::eGFP.

**Figure 1:**
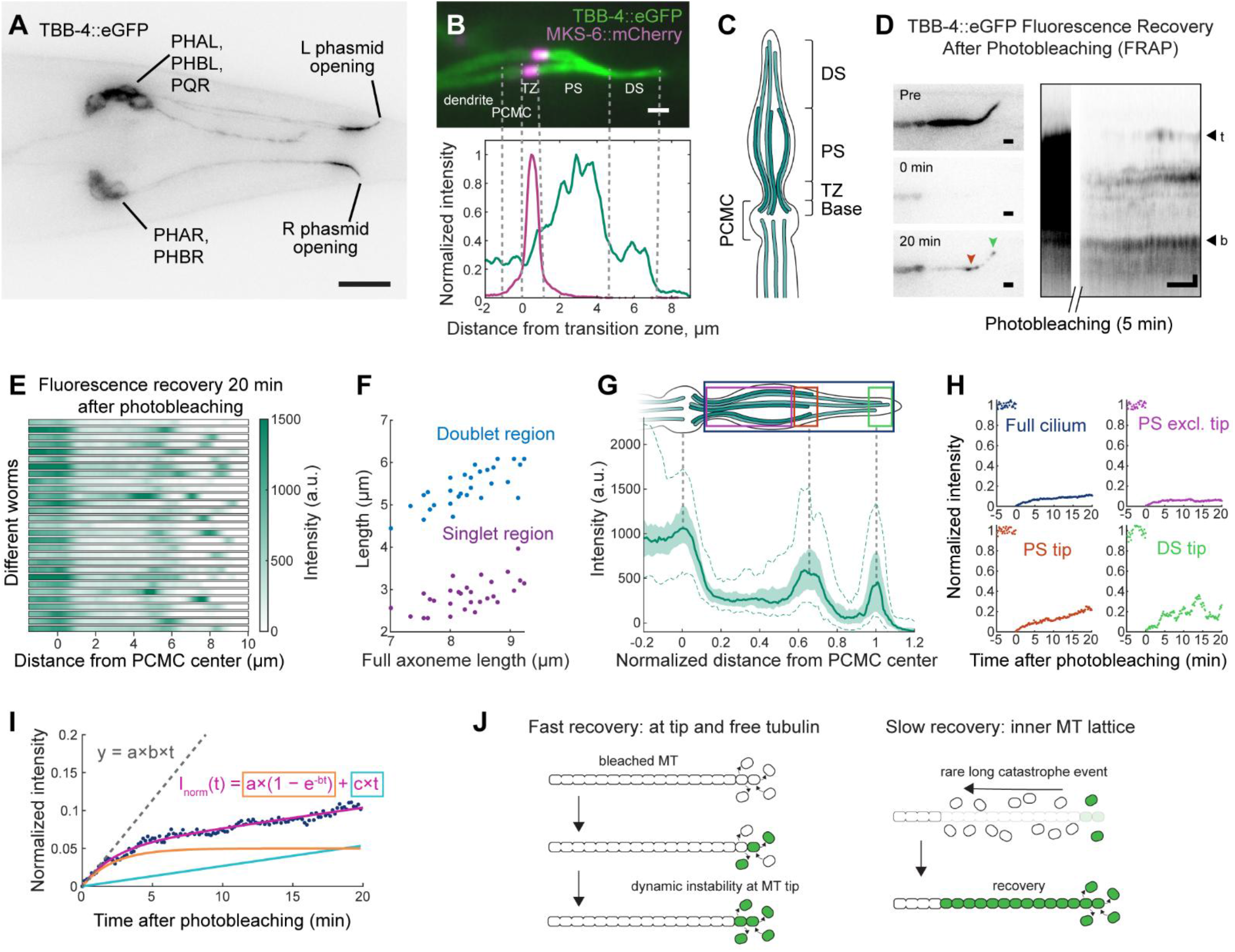
Bulk characterisation of TBB-4 distribution and dynamics in phasmid cilia. **(A)** Inverted fluorescence image of the tail of *C. elegans* expressing TBB-4::eGFP. Scale bar: 10 μm. **(B)** Top: TBB-4::eGFP (green) and MKS-6::mCherry (magenta) localisation in the PHA and PHB cilia. Bottom: TBB-4::eGFP and MKS-6::mCherry intensity profiles normalised to their maxima along the lower cilium in the image above. PCMC, periciliary membrane compartment; TZ, transition zone; PS, proximal segment; DS, distal segment. Scale bar: 1 μm. **(C)** A schematic depiction of axoneme segments indicated in B. Ciliary base, the starting point of axonemal MTs is located before the TZ, within the PCMC. **(D)** Left: Example inverted images of TBB-4::eGFP in the cilia before photobleaching (pre), right after photobleaching (0 min), and after 20 min of fluorescence recovery. Scale bar: 1 μm. Right: a kymograph (time-space intensity plot, intensity-inverted) along the cilium shown on the left, before and after photobleaching. Approximate locations of the ciliary base (b) and tip (t) are indicated. Scale bars: 1 μm (vertical), 5 min (horizontal). **(E)** TBB-4::eGFP intensity distribution along the cilia after 20 min of fluorescence recovery after photobleaching, in n = 29 worms. **(F)** Distance between the PCMC centre and the PS tip peak (doublet region length), and between the PS and DS tip peaks (singlet region length) plotted as a function of distance between the PCMC centre and the DS tip peak (full axoneme length) for worms shown in E. **(G)** TBB-4::eGFP intensity distribution length-normalized to DS tip peak for worms shown in E. Solid line, median; light shading, 25^th^-75^th^ percentiles; dashed lines, 5^th^ and 95^th^ percentiles. A drawing above the plot schematically illustrates the position of the intensity peaks. **(H)** Normalised mean TBB-4::eGFP fluorescence intensity (*I*_*norm*_, see Methods) before and after photobleaching, measured for the cilium shown in D, in the four regions schematically depicted in G (coloured boxes). T = 0 min corresponds to the start/end of photobleaching. Intensity during the photobleaching is not shown. **(I)** Normalised TBB-4::eGFP intensity (*I*_*norm*_) after photobleaching in the full cilium shown in H (blue dots) with a fit *I*_*norm*_(*t*) = *a*(1 − *e*^−*bt*^) + *ct* (magenta line, *a, b, c* – fitting parameters). The exponential (*y* = *a*(1 − *e*^−*bt*^), orange), linear (*y* = *ct*, light blue) components of the equation, and the exponential slope at t = 0 (*y* = *abt*, grey dashed line) are shown. **(J)** A schematic depiction of processes that could underly TBB-4::eGFP fluorescence recovery during the fast (left panel) and slow (right panel) phase. Left: Photobleached tubulins (white) in the free ciliary pool, as well as the ones in the MT lattice close to the plus-tips, are exchanged by the fluorescent tubulins (green) coming from the dendrite. Fluorescence recovery saturates when the percentage of fluorescent tubulin in the most dynamic MT plus-tip regions reaches the prebleached value. Right: Axonemal MTs occasionally shrink further than before, enabling the exchange of tubulin dimers located deeper in the MT lattice.

Intensity profiles of the TBB-4::eGFP fluorescence along the ciliary long axis (Figure 1B) show that TBB-4::eGFP intensity increases from the transition zone, reaching a maximum at 3-4 µm into the cilium, and decreases again into the DS. To unequivocally identify the transition zone, we also imaged MKS-6::mCherry in the same worm. The intrinsic geometry of the PHA/PHB cilia pair results in spatial separation of the proximal segments, whereas the distal segments converge and overlap; consequently, the proximal portions can be resolved when both cilia lie within a single xy focal plane of the microscope (Figure 1B). In other orientations of the worms, the proximal segments might be on top of each other (Supplementary Figure 2A-B). To evaluate how different worm orientations would affect the measured intensity profiles, we simulated two scenarios: complete overlap of the cilia and overlap limited to the distal ~4 µm. In these simulations, we assumed the same axoneme doublet/singlet composition as above and homogeneous incorporation of TBB-4::eGFP in the axoneme. We took into account the resolution of our microscope and that 2.3% of β-tubulin is TBB-4::eGFP. The intensity profiles obtained in this way look similar to the experimental ones (Supplementary Figure 2C-E), with the key difference that the experimental profiles show an intensity increase starting slightly further into the cilium and appearing more gradual than the simulated ones. This indicates that TBB-4::eGFP concentration is substantially lower in the ciliary base and the proximal part of the TZ. A similar observation was previously reported, using an extrachromosomal array–expressed TBB-4 construct [12].

### FRAP experiments show multiscale dynamics at the tips of A- and B-microtubules

To explore the dynamics of TBB-4::eGFP and of axonemal MTs, we first performed fluorescence recovery after photobleaching (FRAP) experiments. We photobleached TBB-4::eGFP in cilia with a cropped 491 nm laser beam at maximum power (~10 W/mm^2^) for 5 minutes and then monitored fluorescence recovery using low-power excitation (~0.25 W/mm^2^) and time-lapse imaging (Supplementary Movie 1). Recovery of fluorescence signals was observed mainly in two locations: the proximal segment (PS) tip and distal segment (DS) tip (Figure 1D-G), similar to a previous study [12]. The positions of the PS and DS tips varied slightly between worms (Figure 1E, F), but the relative distance between PS and DS tips in a single cilium was more constant: the distance from the PCMC centre to the PS tip was ~65 % of the distance from the PCMC centre to the DS tip (Figure 1G). This suggests that most B-tubules cover the first 60-65% of the axonemal length, and the singlet A-tubules – the remaining 30-35%. We also observed additional fluorescent spots appearing within the PS of most cilia (Figure 1E). These spots did not show any consistent localisation across the worms but resulted in the recovered intensity in the PS (excluding the PS tip) being slightly higher than in the DS (Figure 1G). This might be caused either by the presence of a few shorter, dynamic MTs, ‘hotspots’ of tubulin incorporation within the MT lattice, or recovery of the free tubulin pool in the PS.

To further assess the FRAP kinetics, we focused on the time dependence of the integrated fluorescence intensity in four regions: (1) the entire cilium (starting after the PCMC), (2) the PS excluding the tip, (3) the PS tip, and (4) the DS tip (Figure 1G). The integration area was selected slightly larger than the actual fluorescence spots to compensate for small image drift or microtubule instability events occurring during the acquisition; consequently, 100% recovery was not expected. In the PS-excluding-the-tip region, fluorescence intensity reached a plateau within a few minutes after photobleaching, but in the PS-tip and DS-tip regions, no intensity saturation was observed in the first 20 minutes of recovery in most cases (Figure 1H). Often, the DS tip showed rather dramatic intensity fluctuations, with the fluorescence signal almost completely disappearing and reappearing multiple times (Figure 1D and H, Supplementary Figure 3G, Supplementary Movie 2). We next fitted the intensity recovery time traces. In Supplementary Note 1, we show that reliable fits could be obtained to a combination of an exponential function (with a decay time on the order of a minute) with a linear term (Figure 1I). We interpret this linear term as an exponential function decaying too slowly within our measurement window to be fitted accurately, so on at least the tens-of-minutes timescale. We hypothesize that the fast fluorescence recovery is due to a combination of two processes: (1) replacement of photobleached free TBB-4::eGFP in the cilia with fresh, fluorescent protein, arriving from the dendrites, either by IFT or diffusion; and (2) substitution of the photobleached tubulin dimers in the lattice located close to the MT plus-end with fluorescent ones (Figure 1J left panel). The length of axonemal MTs is relatively stable over time, and incorporation of fresh, fluorescent tubulin mostly takes place locally at the PS and DS tips, likely within this fast, minute timescale. Further recovery beyond the MT tips requires replacement of photobleached tubulin located deeper within the MT lattice. This can only occur if MTs depolymerise beyond the section that is replaced with fluorescent tubulin, exposing more proximal, bleached lattice regions. Subsequent rescue then allows incorporation of additional fluorescent tubulin (Figure 1J right panel). Because such long depolymerisation events are rare, the probability of accessing progressively more proximal lattice regions decreases over time. As a result, this process produces a slower, stochastic recovery component, dominated by infrequent episodes of large MT shrinkage followed by rescue.

### TBB-4 diffuses in the dendrites and cilia and is actively transported by anterograde IFT

To get insight into how tubulin is transported from the neuron soma to the cilium, where it is incorporated into MTs, we imaged single TBB-4::eGFP molecules. These experiments were performed using a variant of small-window illumination microscopy (SWIM) [34]. Before acquisition, a region of interest in the PHA/PHB neurons was photobleached by prolonged excitation with a cropped high-intensity 491 nm beam until only a few fluorophores could be detected simultaneously. Next, the sample was imaged continuously, keeping the laser intensity high and adjusting the diameter of the excitation window according to the experimental needs. This approach allowed for minimising the static fluorescence signals arising from TBB-4::eGFP incorporated into the axoneme lattice and enabled long-duration continuous imaging of fresh, fluorescent TBB-4::eGFP proteins entering the bleached region (Figure 2A) [34]. We note that such imaging conditions, in particular the relatively high excitation intensity, caused rapid photobleaching of the majority of eGFP molecules, resulting in single-molecule tracks only lasting, on average, a few seconds. In the following, we first provide a qualitative evaluation of TBB-4 behaviour from kymographs obtained from the image stacks.

**Figure 2:**
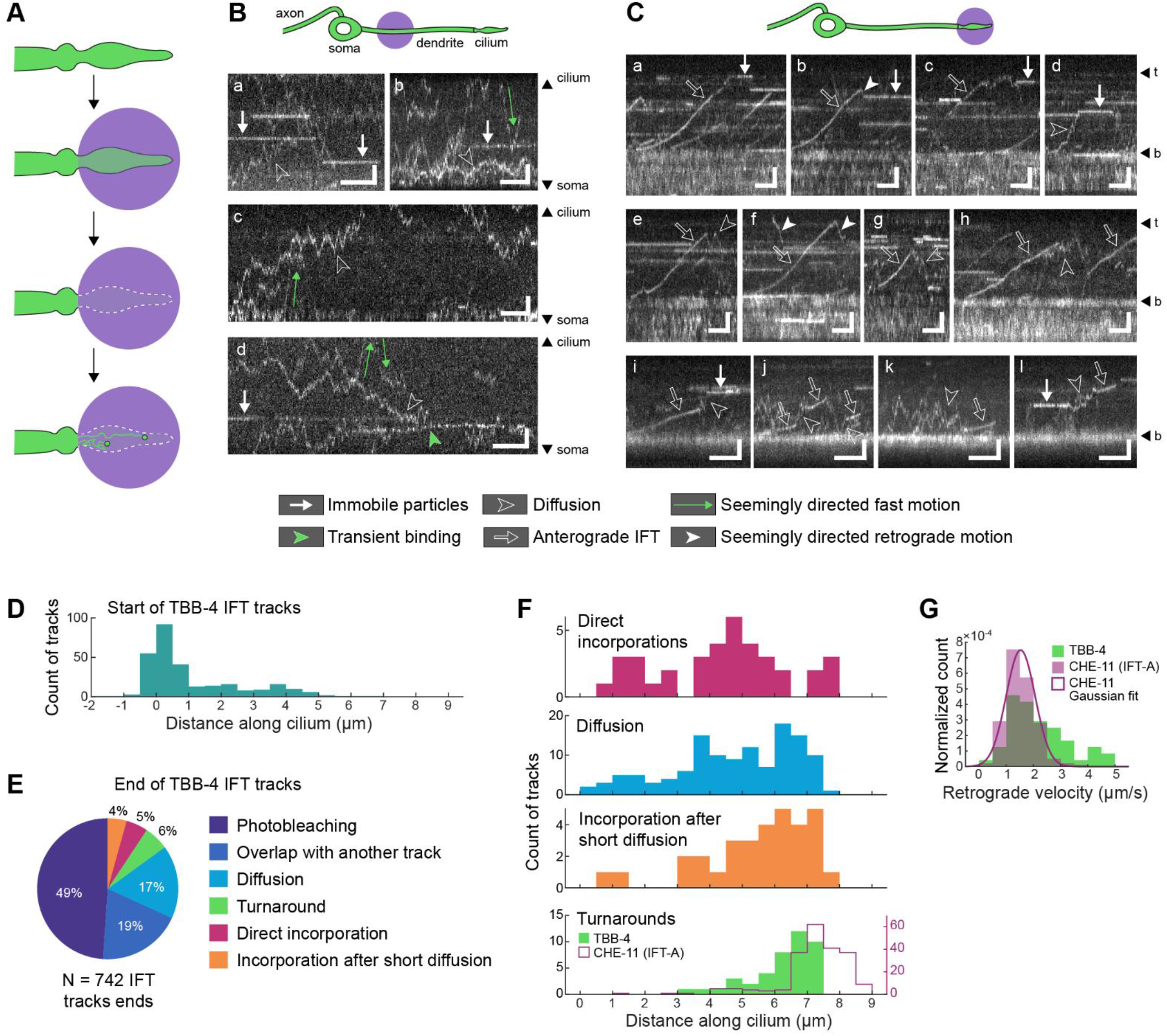
Single-molecule qualitative characterisation of TBB-4 dynamics in phasmid cilia and dendrites. **(A)** A schematic illustration of the imaging approach we used to detect single-molecule TBB-4::eGFP trajectories. A cropped high-intensity 491 nm beam (shown as violet circle) was used to photobleach most fluorophores in the region of interest and then to continuously detect new TBB-4::eGFP entering the bleached area, as described in [34]. **(B)** Representative kymographs showing TBB-4::eGFP dynamics in the dendrites of PHA/PHB neurons, acquired at 37 (b, c) and 46 fps (a, d). Scale bars: 2 µm (vertical), 1 s (horizontal). **(C)** Representative kymographs showing TBB-4::eGFP dynamics in PHA/PHB cilia, acquired at 13 (a-h) and 19 fps (i-l). Approximate positions of ciliary base (b) and tip (t) are indicated. Scale bars: 2 µm (vertical), 2 s (horizontal). Trajectories with different motility characteristics are indicated (manual classification; see legend at the bottom). **(D)** Distribution of starting positions of TBB-4 IFT tracks along the cilium. Also see Supplementary Figure 8B-D. **(E)** Manual classification of the ways TBB-4 IFT tracks end (n = 742 tracks, 38 cilia). See example events for each class in Supplementary Figure 4B. **(F)** Distribution of the ending locations along the cilium for TBB-4 IFT tracks ending with a direct incorporation (magenta), diffusion (blue), brief diffusion followed by an incorporation (orange), or a turnaround (green). The distribution of turnaround locations of an IFT-A component CHE-11 is shown for comparison. As a distance reference point (x = 0), for TBB-4, we used the edge of PCMC, and for CHE-11, the base peak where pausing events occur. These two reference points colocalise (see Figure 3D). Track ends were selected manually on kymographs. **(G)** Velocity distribution for retrograde (apparently) directed movement of TBB-4 (green, 48 tracks) and IFT-A component CHE-11 (purple, 199 tracks). Magenta line, Gaussian fit to the CHE-11 velocity histogram (µ= 1.53, σ = 0.53). Velocities were determined by manually picking two points along a linear-like segment of a retrograde track right after the turnaround (see Supplementary Figure 4C).

In the PHA/PHB dendrites, we mainly observed two kinds of TBB-4 dynamic behaviour: (i) immobile and (ii) diffusive (Figure 2B, Supplementary Movie 3). (i) Immobile particles, visible as horizontal lines in the kymographs, likely correspond to TBB-4 that had been incorporated into the lattice of dendritic MTs. Many of these trajectories appeared ‘out of nowhere’, with the fluorescence abruptly switching on at a random location. This might indicate that these trajectories are due to eGFP blinking ‘on’ again after being photobleached [35]. (ii) Diffusive tracks in the kymographs sometimes showed bouts of apparently confined diffusive motion and bouts where movement seemed more ‘directed’, with velocities of 10-20 µm/s by far exceeding those of the MT motor proteins that drive directed transport in the dendrites [34, 36-40]. We also occasionally observed blinking and slowly moving particles, which might correspond to tubulin molecules transiently binding to growing or shrinking MTs. Only very rarely did we capture directed motion of TBB-4 in the dendrite with velocities close to those expected for cytoplasmic dynein [34] (Supplementary Figure 4A), which could be due to rare events where tubulin is attached to a transport vesicle moving along the dendrite. This qualitative assessment of the kymographs is consistent with the majority of TBB-4::eGFP travelling from soma to cilium by diffusion.

In kymographs of TBB-4::eGFP in the cilia, we observed mainly three types of tubulin dynamics: (i) immobile, (ii) directedly moving in the anterograde direction, and (iii) diffusing (Figure 2C, Supplementary Movies 4 and 5). Like in the dendrites, immobile particles were observed all along the cilia and often appeared ‘out of nowhere’, which could correspond to freshly incorporated fluorescent TBB-4::eGFP, but also to photobleached TBB-4::eGFP already present in the MT lattice, becoming fluorescent again [35]. Occasionally, we observed that immobile particles became diffusive (Figure 2C l), but in most cases, the tracks of immobile particles ended by photobleaching. In some cases, we could see a diffusive or directedly moving particle becoming immobile, likely due to incorporation in the MT lattice (Figure 2C a-d, i). In some directedly moving tracks, immobilisation did not occur instantly but was preceded by a brief inversion of the direction and/or diffusive bout (Figure 2C b, c). In many tracks, we observed bouts of directed motion followed or preceded by diffusion (Figure 2C e, g, h, j-l). This qualitative assessment of the kymographs is consistent with earlier observations showing that both IFT and diffusion contribute to tubulin transport in the cilia [27, 30, 32].

### TBB-4 binds to assembled IFT trains and detaches from them before the trains disassemble

We next proceeded with a more quantitative analysis of the TBB-4 tracks in the kymographs. To this end, we first mapped the starting and ending locations of the anterograde-directed trajectories. As we have shown before [41], directed TBB-4 trajectories start all along the cilium, with a peak at the proximal side of the TZ, adjacent to the PCMC (Figure 2D). This is in contrast to IFT-train components and IFT dynein, which dock exclusively at the ciliary base during train assembly [40-43]. This suggests that tubulin can enter the cilium, at least partially, by diffusion, as has been shown before in *C. reinhardtii* [27, 30]. From the distribution of TBB-4-trajectory starting locations, it is impossible to conclude whether certain factors at the TZ promote TBB-4 binding to anterograde trains or whether this is a purely stochastic process, governed by the balance between tubulin diffusion coefficient and IFT-binding rate. We note that the observed distribution is likely skewed by photobleaching, further complicating the issue (Supplementary Figure 8B-D). It is, however, remarkable that most tracks start on the distal side of the ciliary base, even though assembling trains spend a relatively long time (seconds) at the proximal side of the base [40, 41, 43] and the concentration of soluble tubulin is relatively high there. It might be that during the early stages of IFT train assembly, the IFT-81/74 tubulin binding site is made unavailable to prevent its interaction with axonemal MTs, which would interfere with the proper assembly and positioning of IFT-B complexes at the ciliary base. Possibly, IFT-B proteins undergo certain conformational changes or post-translational modifications at later stages of train assembly, making tubulin binding sites available. In line with this, tubulin was shown to bind IFT trains briefly before their departure from the base in *C. reinhardtii* [43]. However, both these observations are inconsistent with the recently proposed hypothesis that tubulin binds to the IFT81/IFT74 module prior to full anterograde train assembly and subsequently dissociates from the IFT81 CH domain [25], a model proposed to reconcile how tubulin can associate with anterograde trains despite structural data showing that tubulin binding to the IFT81 CH domain sterically clashes with a neighbouring IFT-B repeat [24]. Thus, it remains unclear how tubulin binding to the IFT complex is mechanistically achieved and regulated.

End locations of directed TBB-4 tracks could not be detected in about 70% of cases, either because of photobleaching or because of spatial overlap with other, usually immobile particles (Supplementary Figure 4B top panels). The tracks in which a clear end location of directed transport could be identified were classified into four classes: (i) transitions to the diffusive state, (ii) turnarounds, (iii) direct incorporations, or (iv) incorporations after a short diffusive phase (Figure 2E, Supplementary Figure 4B). Direct incorporations (iii) were observed at the tips of the PS and DS, as well as within the PS, consistent with the FRAP data (Figure 2F, magenta; Figure 1E). We observed more incorporations in the PS than at the DS tip, which is likely caused by photobleaching, making tracks extending into the DS rare. Transitions from directed motion to diffusion ((i) and (iv)) occurred mostly 3-8 µm along the cilia (Figure 2F, blue and orange), which might suggest that the interaction between TBB-4 and IFT trains weakens in the distal part of the cilium. We only very rarely observed directed retrograde TBB-4 tracks. In the majority of cases, this occurred in anterograde tracks turning around (ii), often in the DS. TBB-4 turnarounds appeared instantaneous, without noticeable pause, unlike the IFT-B core component OSM-6, which has been shown to pause at the tip longer than other train components (IFT-A, IFT-dynein, OSM-3) [44]. It is thus likely that TBB-4 is no longer connected to the same IFT-B subcomplex when IFT trains transition from the anterograde to the retrograde state at the ciliary tip. In support of this, we observed that TBB-4 turnarounds occurred on average ~1 µm more proximal than the DS tip (Figure 2F, green and magenta line). Finally, we determined the retrograde velocity in the turnaround tracks of TBB-4 and CHE-11. CHE-11 retrograde velocities were normally distributed with a peak at 1.53 ± 0.074 µm/s (95% CI), consistent with previous studies [45-47]. TBB-4 retrograde velocities, however, were more widely and asymmetrically distributed, with a shoulder extending to velocities as high as 5 µm/s (Figure 2G). We interpret this shoulder to be due to trajectories that appear directed (and are classified like that) but are, in fact, diffusive (see Supplementary Figure 7A and Supplementary Note 3).

Taken together, these results suggest that the binding of TBB-4 to IFT trains is not directly governed by the assembly or disassembly of IFT trains. It appears that TBB-4 only binds to fully assembled anterograde-moving IFT trains [41] and very rarely binds to retrograde trains. The majority of TBB-4 detaches from anterograde trains in the distal half of the cilium, at less specific locations and more proximally than the ciliary tip, where IFT trains turn around [44].

### Binding to the IFT affects TBB-4 distribution

To better understand the interplay between diffusion and IFT-driven transport of tubulin in cilia, we investigated worm strains where putative interactions between the IFT machinery and tubulin had been mutated. It has recently been proposed that DYF-5-mediated phosphorylation of the IFT-74 N-terminal domain (IFT-74N) weakens tubulin-IFT interactions, triggering tubulin unloading [26] (Supplementary Figure 4D). In addition, the reported distribution of DYF-5 along cilia [48] overlaps with the locations where we observed TBB-4 to detach from trains. To test whether IFT-74N phosphorylation affects tubulin transport in cilia, we imaged TBB-4::eGFP in worm strains expressing phospho-dead (PD) IFT-74 (with 11 serine/threonine amino acids substituted by alanine), or phospho-mimic (PM) IFT-74 (with the same amino acids replaced by aspartic and glutamic acids) [26]. We did not observe clear differences in TBB-4 dynamics or distribution between these strains and wild-type control (Supplementary Figure 4E-I). In contrast to an earlier study comparing these *ift-74* mutant strains, the ciliary length covered by TBB-4::eGFP was similar in PD and PM strains (Supplementary Figure 4F). Some worms showed other ciliary defects, such as misalignment of ciliary bases (Supplementary Figure 4E, G). These results suggest that DYF-5-mediated phosphorylation of IFT-74N does not appear to regulate the transport and distribution of tubulin, at least of tubulin-β isotype TBB-4.

We next evaluated the well-characterised interaction between IFT-B protein IFT-74 with the negatively charged C-terminal tail (E-hook) of β-tubulin (Figure 3A) [23, 30, 31]. To weaken this interaction, we generated a *C. elegans* strain ectopically expressing TBB-4::eGFP without its E-hook (ΔE-hook TBB-4), leaving all endogenous tubulin genes unaffected (see Supplementary Note 2). This deletion is analogous to the one used in *C. reinhardtii*, where it was shown to cause a 90% decrease in the frequency of active transport, while incorporation in the axoneme lattice reduced only slightly [30]. In *C. elegans* chemosensory cilia, we observed that while ΔE-hook TBB-4 is present in the somas, dendrites and PCMC at levels similar to those of wild-type TBB-4, its intensity in the cilia is substantially lower (Figure 3B, Supplementary Figure 6B-C). In the cilia, ΔE-hook TBB-4 was detected mainly in the PS, and single-molecule kymographs show that it is almost exclusively diffusive (Figure 3D-F, Supplementary Movie 6). Some of the diffusive tracks appeared more directed (and often faster than expected for IFT). Only very rarely did we observe directed trajectories moving at typical IFT velocities (Figure 3F d, e), ~40x less frequently than in wild type (Figure 3G, Supplementary Figure 6D). Occasionally, we observed diffusive tracks ending with an incorporation (Figure 3F c, d). To confirm that observed differences in TBB-4 distribution and dynamics are not caused by other, non-specific ciliary or IFT defects, we co-expressed both wild-type and ΔE-hook TBB-4::eGFP with an IFT-A marker, CHE-11::mCherry (Figure 3C). In these strains, CHE-11 was distributed similarly along the cilium (Figure 3D) and showed similar signatures of bidirectional active transport (Figure 3E), indicating that ciliary structure and IFT are normal in these strains. Therefore, we can conclude that in the phasmid neurons of *C. elegans*, the absence of the E-hook severely impairs the active transport of TBB-4, resulting in a lower amount of truncated TBB-4 in the cilia.

**Figure 3:**
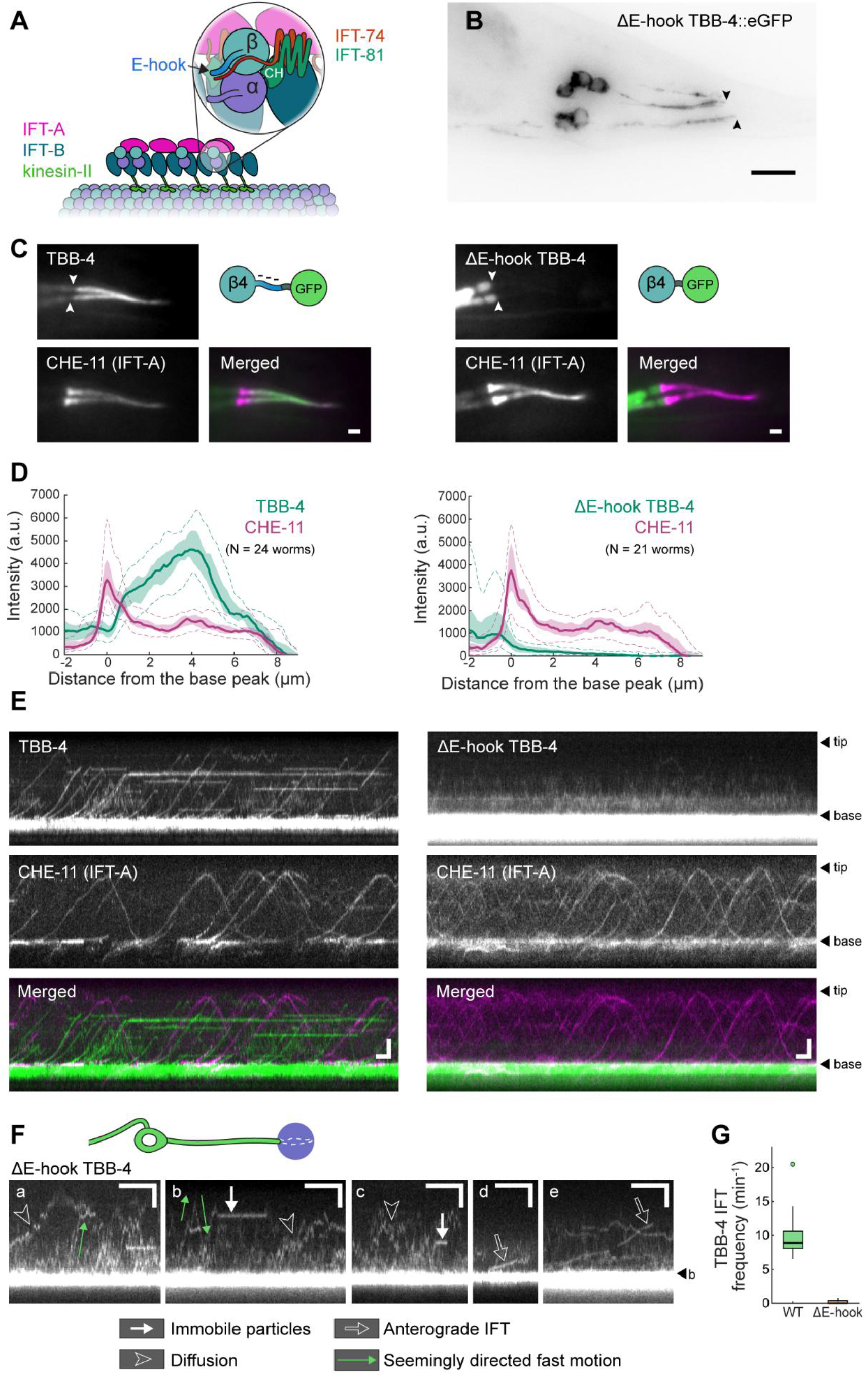
Distribution and dynamics of mutated TBB-4 with impaired connection to IFT trains. **(A)** Schematic depiction of the supposed tubulin binding site on the (anterograde) IFT train. N-terminal domains of IFT-B1 proteins IFT-81 and IFT-74 together bind a tubulin dimer: calponin homology (CH) domain of IFT-81 interacts with globular parts of αβ-tubulin dimer, and the unstructured N-terminal region of IFT-74 forms ion interactions with negatively charged C-terminal tail (CTT) of β-tubulin. Based on [23, 24, 31]. **(B)** Inverted fluorescence microscopy image of the tail of *C. elegans* ectopically expressing ΔE-hook TBB-4::eGFP. The truncated protein is distributed throughout the neurons, similar to WT TBB-4::eGFP (Figure 1A), but is practically absent in the cilia. Arrowheads point at the approximate ciliary base locations (end of PCMC). Scale bar: 10 µm. **(C)** Fluorescence images of WT TBB-4::eGFP (left panel) or ΔE-hook TBB-4::eGFP (right panel) co-expressed with IFT-A component CHE-11::mCherry. Merged images: TBB-4::eGFP, green; CHE-11::mCherry, magenta. Scale bar: 1 µm. **(D)** Intensity profiles of TBB-4::eGFP (WT on the left, ΔE-hook on the right) and CHE-11::mCherry in the cilia. Solid lines, median; light shading, 25^th^-75^th^ percentiles; dashed lines, 5^th^ and 95^th^ percentiles. CHE-11 shows a similar distribution in both strains. ΔE-hook TBB-4 is present in the cilia only in the proximal part and in minimal amounts. Intensity variation in the dendrites and PCMC is higher for ΔE-hook TBB-4 than WT TBB-4, because ΔE-hook TBB-4 was expressed as an extrachromosomal array, so the expression level varied more among the worms (also see Supplementary Figure 6). **(E)** Kymographs showing TBB-4::eGFP (WT on the left, ΔE-hook on the right) and CHE-11::mCherry dynamics in the cilia. CHE-11 undergoes bidirectional active transport in both strains. Unlike WT TBB-4, which is actively transported in the anterograde direction (also see Figure 2C), ΔE-hook TBB-4 is mainly diffusive. Scale bars: 2 µm (vertical), 2 s (horizontal). **(F)** Representative kymographs showing ΔE-hook TBB-4::eGFP dynamics in the cilia in better detail (acquisition rate 19 fps). Short IFT trajectories can be occasionally observed (e, f). Scale bars: 2 µm (vertical), 2 s (horizontal). **(G)** Frequency of anterograde TBB-4 IFT tracks in WT and ΔE-hook TBB-4::eGFP strains. Tracks were manually picked from kymographs if they occurred in the first 1-2 µm of cilia.

### IFT is primarily needed for concentrating tubulin at the tip

The observation that ΔE-hook TBB-4 is largely excluded from the axoneme suggests that purely diffusive tubulin is not able to efficiently reach the sites of its incorporation in the cilia. This could be due to several reasons, which we assess below.

i. Is diffusion much slower than IFT in delivering tubulin to the MT tips? To obtain quantitative insight, we tracked diffusive TBB-4::eGFP particles in the dendrites and cilia (Figure 4A). For the measurements in cilia, we used the ΔE-hook TBB-4 strain, in which TBB-4::eGFP diffuses but is not transported by IFT and exhibits substantially fewer static molecules, thereby facilitating single-molecule diffusion tracking. Most of the diffusive tracks in cilia are located in the PS, close to the central axis of the cilia (Figure 4C). This supports the previous observation that the diffusion of soluble proteins in primary cilia mainly takes place inside the axonemal lumen [32]. From the tracks, we calculated the mean squared displacement (MSD) as a function of time lag and obtained a one-dimensional diffusion coefficient, *D* (Figure 4B) of ~3 µm^2^/s, both in dendrites and cilia. We estimate that tubulin transport by IFT from the ciliary base to the tip takes about 8 seconds (a distance of 8 µm at an average velocity ~1 µm/s). To cover this distance by diffusion, it would take on average 11 seconds (Δ*x*^2^/2*D* ≈ (8 *µm*)^2^/ (2 ∙ (3 *µm*^2^/s))). This is only marginally longer, too small a difference to explain that ΔE-hook deletion results in a severe reduction of TBB-4 incorporation. Moreover, in the dendrites that are substantially longer than cilia (and with the time scaling with the square of distance), diffusion turns out to be a sufficient mode of transportation for tubulin.
ii. Is tubulin unable to efficiently cross the transition zone without IFT? The distribution of localisations where tubulin docks onto IFT trains (Figure 2D) suggests that tubulin can diffusively enter the cilium but does not unequivocally indicate whether IFT plays a substantial role in tubulin transport across the TZ (Supplementary Figure 8B-D). Furthermore, we observed a clear drop in fluorescence intensity between PCMC and cilium in the ΔE-hook TBB-4 strain, which might suggest that TZ acts as a physical barrier for TBB-4. We observed, however, a similar intensity drop for freely diffusing eGFP, which is substantially smaller and not expected to be hampered by the TZ (Figure 4D, Supplementary Figure 7B), which suggests that the observed drops in fluorescence intensity from the PCMC to PS, and further, from PS to DS, reflect the differences in local volume available for free diffusion, rather than protein concentration in these segments (Figure 4E). Thus, our data does not provide strong evidence that IFT is needed for tubulin to enter the cilium, but hints towards it being needed in the distal segment, where the space available for diffusion becomes too limited to supply a sufficient amount of tubulin to the tip.
iii. Is IFT required to maintain an optimal tubulin concentration near the MT plus-tip where soluble GTP-tubulin is incorporated in the tubulin lattice? To test how a combination of directed anterograde transport and diffusion affects TBB-4 ciliary concentration profiles, we performed computer simulations (Figure 4F, Supplementary Figure 8A). Here, we modelled diffusion as a one-dimensional random walk characterised by a diffusion coefficient, *D*, starting at the base of the 8 μm-long cilium. Every diffusive molecule has a probability to bind to an IFT train, characterised by the binding rate parameter *λ*_*bind*_ (for a more detailed description, see Methods). Once bound to an IFT train, the particle remains bound until it reaches the tip, where it becomes diffusive again. These simulations show that in the absence of IFT (*λ*_*bind*_ = 0), tubulin concentration is constant along the cilium. Increasing *λ*_*bind*_ by even a small amount results in a steep tubulin gradient toward the ciliary tip, directly supporting a central role for IFT in concentrating soluble tubulin at incorporation sites (Figure 4G, Supplementary Figure 8H).

Taken together, our experiments and simulations suggest that the main role of IFT in establishing axonemal composition and length is to create a tubulin concentration gradient within the cilium, peaking at the ciliary tips. This regulation of tubulin concentration along the cilium might be crucial to maintain the MT tip dynamics, and subtle changes in the IFT characteristics, such as tubulin binding rate, would significantly alter the tip dynamics.

**Figure 4:**
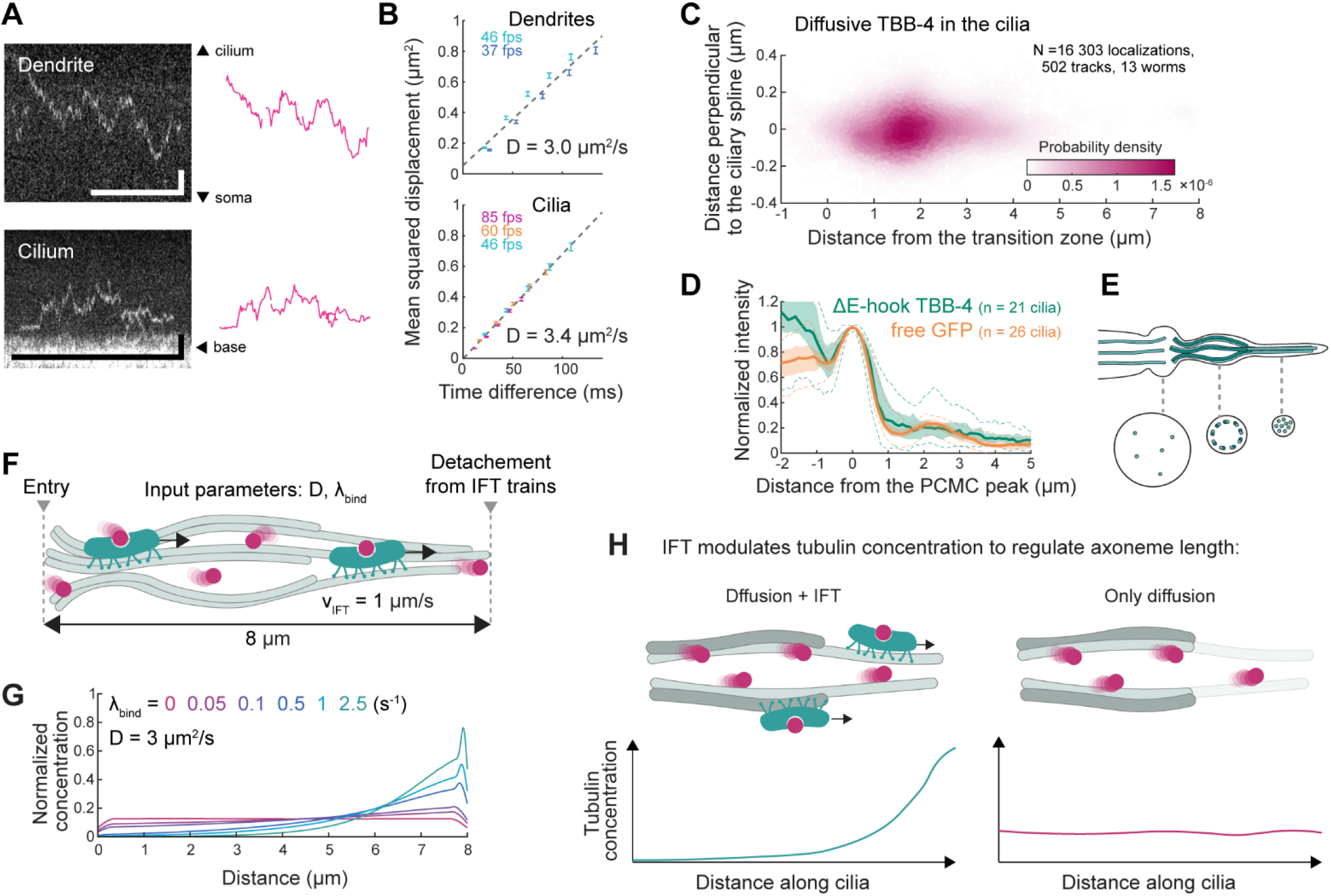
Quantifying TBB-4::eGFP diffusion. **(A)** Left: kymograph fragments showing TBB-4::eGFP diffusion in a dendrite (top) and in a cilium (bottom). Scale bars: 2 µm (vertical), 2 s (horizontal). Right: trajectories obtained from these fragments (magenta lines). **(B)** Mean squared displacement (MSD) parallel to the ciliary spline plotted as a function of time for diffusive tracks in the dendrites (top) and cilia (bottom). One-dimensional diffusion coefficient *D* was extracted by fitting the first 5 data points as *MSD*(*t*) = 2*Dt* + *b* (*b* is an offset arising from experimental noise [49]). **(C)** Distribution of diffusive trajectories in the cilia. The highest density of diffusive localisations is found at the start of the PS, along the central axis of the cilium. **(D)** Intensity profiles along cilia for ΔE-hook TBB-4::eGFP (green) and eGFP (orange). To account for the differences in expression levels between strains/worms, individual plots were normalised to the PCMC peak. Solid line, median; light shading, 25^th^-75^th^ percentiles; dashed lines, 5^th^ and 95^th^ percentiles. **(E)** A schematic illustration explaining the intensity differences observed in D. While fluorophore concentration can be the same, volumes available for diffusion decrease from the dendrite to the cilium, and within the cilium, from PS to DS. **(F)** A cartoon illustrating the simulation used to mimic tubulin dynamics in the cilia (see Methods for more details). Particles start in a diffusive state (with one-dimensional diffusion coefficient *D*) at the ciliary base (*x* = 0). At every time step *t*_*i*_, every particle can either remain diffusive or bind to an anterograde IFT train moving with a velocity of 1 µm/s. Equal probability to bind to an IFT train (*p*) at every time step results in the exponential distribution of IFT-bound particles over time, characterised by binding rate parameter *λ*_*bind*_ (see Supplementary Figure 8A). Once bound to an IFT train, particles remain bound until they reach the tip (*x* = 8 μm), where they become diffusive again. **(G)** Distribution of particles along the ‘cilium’ 500 s after the start of the simulation. A higher IFT-binding rate creates a more prominent concentration peak at the ‘tip’. See example simulated tracks in Supplementary Figure 8H. **(H)** A proposed model of axoneme length and composition regulation via directed transport of tubulin: anterograde IFT concentrates tubulin at the tips, promoting axonemal MT stability (left panel). In the absence of IFT, diffusion-driven tubulin concentration is the same throughout the cilium, meaning fewer molecules in the narrow distal segment, which causes depolymerisation of axonemal MTs, especially the singlet ones (right panel).

## Discussion

Using FRAP and single-molecule fluorescence imaging of β-tubulin isotype TBB-4 in phasmid neurons of *C. elegans*, we have shown that plus-tips of axonemal MTs undergo continuous tubulin exchange, with new tubulin units at least partially arriving from outside of the cilium (Figure 1). Tubulin diffuses from the soma to the ciliary base across a ~50 µm-long dendrite, while within the cilium, transport modes include both diffusion and anterograde IFT (Figure 2B-C). Diffusion rates in both regions are similar, *D* ≈ 3 µm^2^/s (Figure 4B). Disrupting TBB-4’s ability to bind to an IFT train, by removing its C-terminal E-hook, leads to a drastic decrease in its share in the axoneme (Figure 3, Supplementary Figure 6C).

Axoneme dynamics and tubulin transport have been studied extensively in *C. reinhardtii*, the motile cilia of which can be optically separated from the cell body using total internal reflection fluorescence (TIRF) microscopy, and for which flagellar regeneration can be induced in a controlled manner [27, 30, 31, 50]. Similar studies in primary cilia have been lacking, most likely due to experimental challenges: in cell cultures and multicellular organisms, the orientation of cilia relative to the imaging plane is more difficult to control than in *C. reinhardtii*, and surrounding cells create an autofluorescence background, in addition to the static signal arising from the axoneme lattice-incorporated tubulin. It has been suggested that active transport of tubulin is downregulated in steady-state primary cilia, similar to full-length flagella, explaining the observed faint tubulin tracks [12, 51]. Here, by exciting and continuously photobleaching just a small region in the sample, we managed to observe anterograde IFT and diffusion of TBB-4 in the cilia and dendrites of phasmid chemosensory neurons in *C. elegans*. Our approach allowed us to eliminate the static pool of fluorescent tubulin and facilitated the detection of less abundant, moving particles entering the photobleached area, as well as enhancing the signal-to-background ratio by reducing out-of-focus fluorescence [34].

We observed active, IFT-driven transport of TBB-4 in steady-state phasmid cilia at a rate of ~10 tracks/min (Figure 3G, Supplementary Figure 6D). The tubulin dimer influx rate is estimated to be ~7–15 s^−1^, assuming a labelling density of ~2.3% (Supplementary Figure 1E), that each tracked event corresponds to one to two TBB-4::eGFP molecules, and that fluorescent labelling does not alter TBB-4 affinity for IFT. This is comparable to the ~4-8 s^-1^ estimated for fast regenerating *Chlamydomonas* flagella, but much higher than the ~0.3-0.6 s^-1^ for steady-state, fully grown flagella [30]. We note, however, that *C. reinhardtii* expresses only one α-tubulin and one β-tubulin isotype, while *C. elegans* has 9 α-, 6 β- and 1 γ-tubulins. The influx rates of other isotypes might differ from the values we estimated from TBB-4.

Further evidence that active tubulin transport in *C. elegans* chemosensory cilia in steady-state cilia is similar to that of fast regenerating *C. reinhardtii* flagella comes from our FRAP data, from which we can estimate the rate of tubulin dimer ciliary entry to be ~5 × 10^3^ min^-1^ (based on the fluorescence recovery rate *ab*, Supplementary Table 2, and total number of tubulins in the cilia, Supplementary Figure 1B), comparable to ~9 × 10^3^ min^-1^ in fast regenerating *Chlamydomonas* flagella [28]. Despite such active ciliary entry and anterograde transport of tubulin, we observed that the length of the axonemes in *C. elegans* chemosensory cilia is relatively stable, and only microtubule tips appear to undergo constant tubulin exchange, implying frequent catastrophe and rescue events (Figure 1E, [12]). It remains an open question whether tubulin transport in these chemosensory neurons is further upregulated during ciliary growth. We note that, based on the estimated tubulin influx rate, both in *C. elegans* and *C. reinhardtii*, IFT can only provide a fraction of the total required tubulin amount: 3-5% of tubulin needed for the fast phase of axonemal growth in *C. reinhardtii*, and 8-18% of fresh tubulin required for turnover at the tips in *C. elegans*. Consequently, it is likely that the majority of tubulin is supplied by diffusion, as has been suggested for *C. reinhardtii* [30].

In line with these differences in tubulin transport activity, we also discovered that binding to IFT trains has a different impact on tubulin presence in the axoneme of *C. elegans* compared to *C. reinhardtii*. In flagella, removing the E-hook of β-tubulin resulted in 90% less binding to IFT trains, while its presence in the axoneme decreased only by 17% [30]. Moreover, *Chlamydomonas* strains with either mutated CH domains of IFT81 (*ift81-1 IFT81(5E)*) or truncated IFT74 N-terminal domains (*ift72-2 IFT74Δ130*), in which the frequency of tubulin transport was reduced by 74% and 89%, respectively, could nevertheless form nearly full-length flagella, albeit at reduced rates (64% and 78% reduction compared to wild type) [31]. In contrast, in our system, ΔE-hook TBB-4, with impaired ability to bind to IFT trains, was barely present in the axoneme (both, frequency of directed tubulin transport as well as its intensity in the cilia were reduced by ~95%, compared to wild type, Figure 3G, Supplementary Figure 6C-D).

We explored several possible explanations for why binding to IFT is so crucial for tubulin localisation in the axoneme. The speed factor does not seem to play a role, since with *D* ≈ 3 µm^2^/s and only ~8 μm length to cover, diffusion is almost as fast as IFT. For larger proteins, with substantially lower diffusion coefficients (*D* < 1 µm^2^/s), the difference in transport time between diffusion and IFT may become substantial and crucial. In such a case, even intermittent transport by IFT followed by diffusion could substantially decrease the time needed to cross the cilium (Supplementary Figure 8G), as we have observed before for the ciliary membrane protein OCR-2 [52]. It has been shown that small proteins can enter cilia by diffusion, while larger ones rely on active transport through the ciliary gate because the sieving mesh size of the TZ is too small for them [53-56]. eGFP-labelled tubulin dimers might be on the edge, as their molecular weight exceeds this limit, but they are not a single globular protein. Further, the observations that the ΔE-hook TBB-4::eGFP signal is distributed similarly to that of free eGFP (Figure 4D), and that IFT docking locations of TBB-4 extend into the cilium (Figure 2D), let us conclude that tubulin does not require IFT for entering the cilium. Finally, using computer simulations, we assessed how a combination of anterograde IFT and diffusion affects the distribution of tubulin inside the cilium (Figure 4G). Since MT dynamic instability takes place at the tips, it is the local concentration of tubulin around the MT tips that matters. In this case, a high diffusion coefficient acts against concentrating tubulin at the tips, as freely diffusing tubulin would quickly spread along the cilium, unless something captures it in place. Binding to the IFT trains is then needed to promote a higher local concentration of soluble tubulin near the MT tips (Figure 4H). A similar explanation has been suggested for the IFT role in regenerating flagella of *C. reinhardtii* [27, 30]. In the chemosensory cilia of *C. elegans*, however, a similar degree of ciliary entry and anterograde transport of tubulin is required just to maintain steady-state axonemes. This might be due to relatively long, more dynamic singlet regions of the MTs in *C. elegans* [11, 57], as well as different MAPs and MIPs present in these cilia, affecting MT stability [6, 10, 11]. In line with this hypothesis, we previously demonstrated that redistribution of IFT to the ciliary base, induced by chemical stimulation [58] or by dendritic ablation [59], effectively switches off IFT-driven tubulin transport and leads to a rapid collapse of the axonemal DS. Besides that, *C. reinhardtii* flagella have been demonstrated to concentrate proteins, including tubulin, at the tips by reducing their diffusivity, possibly via interactions with other tip region proteins [27, 50], which we have not observed in *C. elegans*. Furthermore, a stronger need for IFT might be explained by the geometry of phasmid cilia of *C. elegans*: they bulge in the PS and narrow down in the DS, leaving less volume available for proteins to diffuse (Figure 4D-E, [33, 52]) and making closely located MT tips compete for the same pool of tubulin, thereby decreasing its effective concentration.

Maintaining a steady-state axoneme by constantly supplying tubulin by IFT might have an evolutionary value related to the sensory function of these cilia, despite being energetically more costly. In *C. elegans*, only the tips of the cilia are exposed to the external environment, and the specific location and composition of these regions determine the worm’s ability to sense, respond to stimuli, and ultimately to survive. An axoneme that can be dynamically tuned by regulating the tubulin-loading of IFT trains might therefore be useful in adapting the sensory range of the cilium to the changing environment.

In summary, our single-molecule imaging approach has revealed the dynamics of β-tubulin isotype TBB-4 in phasmid cilia and dendrites of *C. elegans*, demonstrating that IFT-driven tubulin transport is active and essential for maintaining steady-state cilia. Whether this reflects a fundamental distinction between primary and motile cilia, or a feature unique to this organism or cilia type, will require further investigation. The mechanisms regulating tubulin loading and unloading – both in steady-state and growing cilia – also remain to be elucidated. Further research in diverse model systems will be essential to address these open questions.

## Materials and Methods

### C. elegans strains

Strains used in this study are listed in Supplementary Table 1. Worms were maintained according to standard procedures [60], at 20°C, on nematode growth medium (NGM) plates seeded with HB101 *E. coli*. TBB-4::eGFP and CHE-11::mCherry strains were previously generated in our lab using Mos-1-mediated single-copy insertion [61]. To generate extrachromosomal arrays, the same Ptbb-4::tbb-4::egfp::3’UTR(tbb-4) construct as used in the MosSCI strain was cloned into pCFJ350 plasmid (Addgene) and co-injected together with a myo-2::mCherry injection marker. The ΔE-hook construct was cloned by Gibson assembly using two fragments: the gel-purified plasmid backbone with Ptbb-4::tbb-4 cut out, and Ptbb-4::tbb-4 without E-hook (last 42 bp) amplified by PCR. The resulting clones were verified using restriction analysis and sequencing. The details on the construct sequence and primers used can be found in Supplementary Table 3.

### Fluorescence microscopy

#### Setup

Fluorescence data were acquired using a custom-built epi-illuminated widefield microscope [45, 62]. In short, the setup was built around an inverted microscope body (Nikon Ti E), equipped with a 100x objective (Nikon, CFI Apo TIRF 100x, NA 1.49). The system was controlled using MicroManager software (v1.4) [63]. 488 and 561 nm DPSS lasers (Cobolt Calypso and Cobolt Jive, 50 mW) were used for fluorescence excitation, excitation power was adjusted using an acousto-optic tuneable filter (AOTF, AA Optoelectronics) and a neutral density filter (Thorlabs, ND10B) according to the needs of the experiment (see below). Fluorescence emitted by eGFP and/or mCherry was collected by the same objective, filtered from the excitation light using a dichroic mirror (ZT 405/488/561rpc, Chroma), and then from each other using a set of dichroic (565LP) and emission filters (525/45, Brightline HC, Semrock for eGFP; 630/92 for mCherry) inside an image splitter (Optosplit III, Cairn Research). Signal from one or both channels was projected on the EMCCD camera (Andor iXon 897). To excite and photobleach only a small region of interest, the excitation beam was cropped using an iris diaphragm (Thorlabs, SM1D12, ø0.8-12 mm) as described in [34], resulting in a narrow excitation window.

#### Sample preparation

Young adult worms with 6-10 eggs were anaesthetised in a droplet of 5 mM levamisole in M9, sandwiched between a coverslip and an agarose pad (2% agarose in M9), sealed with melted VaLaP (1:1:1 vaseline, lanolin, paraffin) to prevent dehydration, and mounted on the microscope. More details on the sample preparation are provided in [62].

#### Imaging settings

Imaging parameters, such as the size of the excitation window, excitation intensity, and frame rate, were adjusted depending on the concentration of labelled protein, its rate of entering the ROI and the motility one attempts to capture. For bulk measurements (FRAP, intensity profiles), the acquisition was performed using ~2% of the maximal power (~0.2 W/mm^2^ at the centre of the beam), with a 1-5 s interval between the frames to minimise photobleaching. To get rid of the static TBB-4::eGFP signal present in the microtubules, the region of interest was photobleached with the maximal possible 491 nm laser intensity (~10 W/mm^2^ at the centre of the beam), typically for ~5 min for cilia. Consequent imaging at 6.7 fps allowed capturing new static and IFT-transported molecules, while diffusive motion stayed mostly unresolved. This relatively low frame rate allowed using lower laser power (~40% of the maximum), thereby reducing photobleaching and obtaining longer trajectories. To avoid too many fluorescent particles from accumulating in the cilia, the size of the excitation window was made larger (up to 30 µm). Capturing diffusive motion worked best in our hands at 20-100 fps. Due to short exposure time, the excitation intensity needed to be higher (100% of the maximum), and the beam size smaller, to avoid bleaching too many fluorophores at once. Intermediate values were used as well, for example, 13.3 fps at maximal laser power with various excitation window widths. When imaging eGFP and mCherry simultaneously, an alternating excitation strategy was used, as described by Zhang et al. [64], to minimise mCherry photobleaching and detection of fluorescence excited by 488 nm in the mCherry channel.

### Data analysis

#### Overlapping images from two spectral channels

To correct for the x-y shift between two detection channels for eGFP and mCherry imaged in the same sample, we made images of fluorescent beads (TetraSpeck™ 0.1 µm, Thermo Fisher) visible in both channels and aligned them using a Descriptor-based registration (2d/3d) Fiji plugin (affine model). The same model was then applied to the *C. elegans* data.

#### Fluorescence intensity analysis

Intensity profiles were measured along a segmented spline drawn through the centre of a cilium on a time-averaged projection of 10 consecutive frames, using a Plot Profile Fiji function (line width = 7 pixels). This way, the fluorescence intensity is, in effect, integrated along the two axes perpendicular to the ciliary long axis (within the image plane by summing pixels, perpendicular to it by our imaging approach). For background correction, each line was first drawn over an area in the worm close to the measured region but without FP expression, and an average of the first 30 pixels was subtracted from all the intensity values. Integrated or mean intensity within a specific region of interest was measured using a Measure/Measure stack Fiji function. For background correction, the mean intensity of an area close to the ROI without FP expression was subtracted from the measured values.

The profiles of individual cilia are presented normalised to their maximum. Data from multiple cilia are presented as median, 25^th^-75^th^ and 5^th^-95^th^ percentiles of values from individual cilia imaged using the same conditions, without prior intensity normalisation.

#### Ciliary length analysis

To determine the ciliary length, a spline was drawn on a time-average projection of 10 consecutive frames, from the end of the PCMC to the ciliary tip. Since PHA and PHB cilia often overlap in the distal region, the tips of the two cilia often cannot be resolved. The length difference Δ*L* therefore reflects the misalignment of the ciliary bases with respect to the ciliary tip, as illustrated in Supplementary Figure 4G.

#### FRAP data fitting

For each ROI, the mean intensity over time was corrected for background by subtracting the mean intensity of a corresponding background region: *I*(*t*) = *I*_*ROI*_*Mean*_(*t*) − *I*_*BG*_*Mean*_(*t*). Then it was normalised by subtracting the first value after photobleaching (*I*_0_) and dividing by the mean prebleached value (*I*_*pre*_): *I*_*norm*_(*t*) = (*I*(*t*) − *I*_0_)/*I*_*pre*_ . The obtained FRAP curve was fitted with a single-exponential equation (*I*_*norm*_(*t*) = *a*(1 − *e*^−*bt*^)), a combination of an exponential and a linear term (*I*_*norm*_(*t*) = *a*(1 − *e*^−*bt*^) + *ct*), or a double-exponential equation (*I*_*norm*_(*t*) = *a*(1 − *e*^−*bt*^) + *c*(1 − *e*^−*dt*^)), using the nonlinear least-squares method.

#### Kymograph generation

Kymographs were generated along a hand-drawn segmented spline (width = 5 pixels), based on a time-averaged projection of the stack. For initial dynamics visualisation, we used a Multi Kymograph Fiji plugin, for creating high-quality kymographs used in the figures – KymographClear [65].

#### Single-particle tracking

TBB-4 molecules were tracked using a MATLAB-based software, FIESTA (version 1.6.0) [66]. Tracking parameters were adjusted depending on what kind of motility was expected to be captured. Obtained tracks contain information regarding time *t, x* and *y* coordinates, and the distance moved for every tracked time frame. Tracks were visualised as an overlay with a kymograph, and erroneous or very short tracks (<15 frames) were excluded from further analysis. Erroneous tracks occurred when two or more particles were too close to each other, or in the proximity of a bright unbleached region (PCMC, dendrite).

To allow for comparing the data obtained in different worms, the *x*-*y* track coordinates were transformed to the cilia/dendrite-based coordinates, using a MATLAB script. This was done by defining a spline along the cilium/dendrite (by manually drawing a segmented line on an average or maximum projection of a frame stack and interpolating it with a cubic spline curve) and a reference *x* = 0 point (in most cases, the end of the PCMC; manually selected on an average or maximum stack projection), and obtaining distance parallel and perpendicular to the spline for every tracked point (*d*_||_(*t*), *d*_⊥_(*t*), Supplementary Figure 5A) .

#### MSD-based filtering

For each datapoint (*d*_||_(*t*), *d*_⊥_(*t*)), a windowed Mean Squared Displacement classifier (wMSDc) approach, described in Danné et al. [67], was used to extract the anomalous exponent value (α) of the time lag (τ) from *MSD* = 2Γτ^*α*^ (where Γ is the generalised transport coefficient), in the direction parallel (*α*_||_(*t*)) and perpendicular (*α*_⊥_(*t*)) to the spline. *α* was calculated analytically, using the following equation: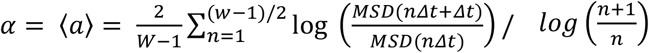 keeping a fixed window *W* = 15 time frames. In this paper, *α*_⊥_ values are referred to as *α*, since we primarily discuss the motion along the length of cilia and dendrites. Obtained *α* values were used to separate trajectories belonging to immobile (*α* ≈ 0) and transported in a directed manner (*α* ≈ 2) TBB-4. Points with *α* < 1 were classified as immobile, and with *α* > 1.2 as IFT-bound (Supplementary Figure 5B-C). This classification doesn’t take into consideration diffusive or temporally pausing particles because such events either were not tracked or were manually excluded from the analysis.

#### Manual selection of events

Events that could not be reliably extracted from single-particle trajectories, such as various kinds of TBB-4 IFT track ends and retrograde-directed motion, were manually picked on the kymographs and saved using a MATLAB script. The intensity border between the PCMC and the TZ, clearly visible on kymographs, was selected as a distance reference (distance along cilium = 0). Retrograde velocities were measured as *v* = *distance*/*time* on linear parts of tracks following the turnaround, as illustrated in Supplementary Figure 4C.

#### Diffusion analysis

Displacement along the length of the cilium or dendrite was calculated for discrete time points for all diffusive tracks. The displacement data from datasets with different frame rates were cumulated to calculate the average mean squared displacement (MSD) for every discrete time point. The first five points of the MSD from each dataset were first fitted with a power equation *MSD*(*τ*) = 2*Γ*τ^*α*^ to estimate whether, under this imaging condition, we captured free or restricted diffusion. MSD values with α > 0.8 were pooled together to calculate the average (1d) diffusion coefficient as the slope of the polynomial fit (*MSD*(*τ*) = 2*Dτ* + *c*) through the first five points.

### Computer simulations

#### TBB-4 intensity profile along one cilium depending on the orientation of the other cilium

Intensity profiles were simulated for each cilium individually, assuming an axoneme to have a total length of 8 µm, of which 5 µm are doublet and 3 µm – singlet MTs. Lattice step was 8 nm (~tubulin dimer size). Each lattice point can accommodate one fluorophore, resulting in an intensity distribution described by a Gaussian function with FWHM = 0.5 μm. For each lattice point, intensity was determined as a random value from a normal distribution (µ= 0.023, σ = 0.005 – estimated labelling density, assuming TBB-4::eGFP being incorporated homogenously throughout the axonemal MTs) multiplied by 9*13 (number of protofilaments in 9 A-tubules, for the singlet region) or 9*(10+13) (A+B tubules, for the doublet region). The region where two cilia start to overlap could be adjusted between 0 to 1 (0 – only one cilium contributes to the sum intensity, 0.2 – the most distal 20% of the second cilium contributes to the sum intensity, etc.).

#### Diffusion with and without IFT

Diffusion was simulated as a 1D random walk. IFT was simulated as unidirectional movement with *v* = 1 μm/s (for simplification). The probability of a diffusive molecule binding to an IFT train was determined by *p* per Δ*t*, which, in the case of many molecules, translates into a binding rate *λ*_*bind*_. All molecules start at the base (*x* = 0). *t*_*tip*_ is the time it takes for a molecule to reach the ‘tip’ (*x* = 8 μm). Once bound to an IFT train, the particle remains bound until reaching the tip, where it dissociates and becomes diffusive again. At the base and at the tip, diffusive particles are elastically reflected. When considering the photobleaching effect, the molecules were set to disappear following an exponential distribution with a characteristic time *t*_*bleach*_ = 1.6s, determined experimentally earlier [41].

#### Estimating the average value and error for distributions

The bootstrapping method was used to estimate the parameters of any given distribution. A distribution consisting of N measurements was randomly resampled (with replacement), and a median of the resampled distribution was calculated. This procedure was repeated 1000 times, resulting in a bootstrapping distribution of medians. The mean (μ) and standard deviation (σ) of this distribution were used to estimate the parameter and its error. All values and errors in this paper are presented as μ ± 3σ.

## Data availability

The experimental data and analysis scripts underlying this study are available on DataverseNL: https://doi.org/10.34894/QZEJPZ

## Author contributions

Conceptualisation: E.L., A.M. and E.J.G.P; data curation: E.L. and A.M.; formal analysis: E. L. and D.G.; funding acquisition: E.J.G.P. and A.M.; investigation: E.L. and D.G.; methodology: E.L., A.M. and E.J.G.P.; project administration: E.L. and E.J.G.P.; resources: E.L.; software: E.L. and A.M.; supervision: E.J.G.P.; validation: E.L.; visualisation: E.L. and A.M.; writing – original draft: E.L.; writing – review and editing: E.L., A.M., D.G. and E.J.G.P.

## Acknowledgements

We acknowledge financial support from the European Research Council under the European Union’s Horizon 2020 research and innovation programme (Grant agreement no. 788363; “HITSCIL”; E.J.G.P.), Marie Sklodowska-Curie Actions Postdoctoral Fellowship of the European Commission (Project no. 898006; ‘MingleIFT’, A.M.) and Czech Science Foundation (Grant no. 26-22371M, A.M.)

