## Supplementary Information for "Intraflagellar transport of tubulin maintains steady-state axoneme integrity in *C. elegans* cilia"

for the article "**Intraflagellar transport of tubulin maintains steady-state axoneme integrity in *C. elegans* cilia**"


### Supplementary Figures

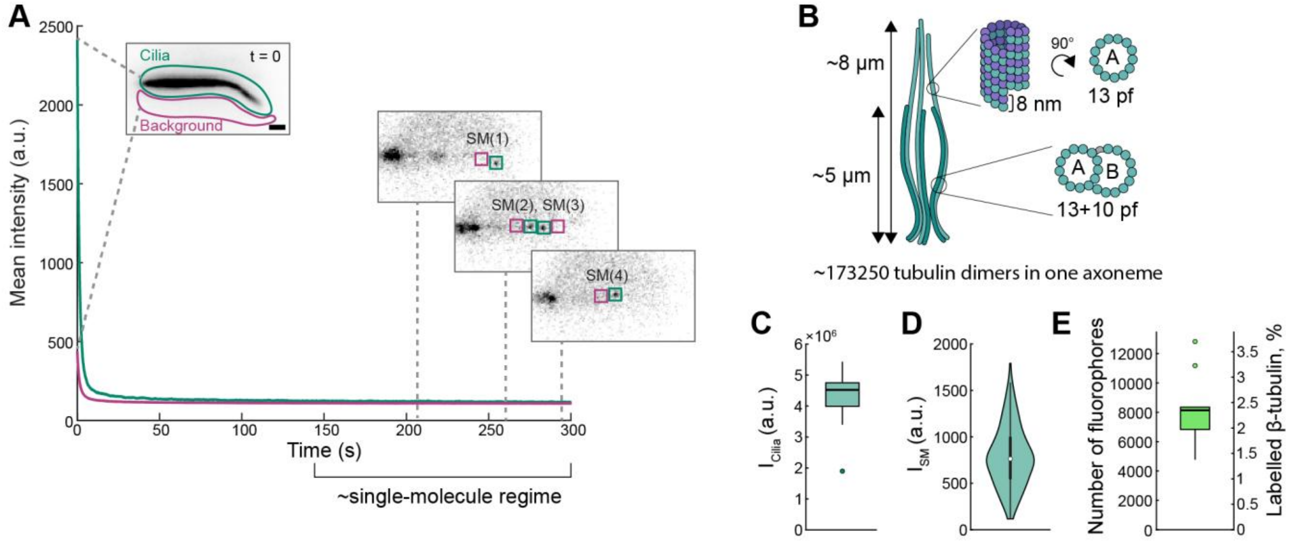


**Supplementary Figure 1: Estimating TBB-4::eGFP amount in the cilia. (A)** TBB-4::eGFP in the cilia (inverted image shown) is photobleached using high-intensity cropped 491 nm beam for 5 min to enable the detection of single eGFP. The mean fluorescence intensity changes over time in the regions containing a pair of cilia ($I_{Mean_{Cilia}}$, green line) and background ($I_{Mean_{Background}}$, magenta line) are plotted. Scale bar: 1 µm. **(B)** Estimating the total number of tubulin dimers in an axoneme comprised of 9 MT doublets, each consisting of an 8 µm-long A-tubule (13 protofilaments, pf) and a 5 µm-long B-tubule (10 pf). Given the length of αβ-tubulin dimer is ~8 nm, one axoneme contains 173250 tubulin dimers. **(C)** Integrated intensity in unbleached cilia calculated as $I_{Cilia} = (I_{Mean_{Cilia}}(t=0) - I_{Mean_{Background}}(t=0)) \times Area_{Cilia}$ (n = 11 cilia pairs/worms). **(D)** Violin plot of single eGFP intensities calculated as $I_{SM} = (I_{Mean_{SM}} - I_{Mean_{Background}}) \times Area_{SM}$ (n = 379 fluorophores, 11 cilia pairs/worms). Examples of single eGFP intensity measurements are shown in A (green box, single eGFP; magenta box, corresponding background). **(E)** Left axis: The number of TBB-4::eGFP in a pair of cilia estimated as $N_{fluorophores} = I_{Cilia}/Mean(I_{SM})$ (n = 11 cilia pairs/worms). Right: The percentage of labelled tubulin in the cilia estimated as $N_{fluorophores}/2 \times 173250\ (n_{tubulin\ dimers\ per\ axoneme})$.

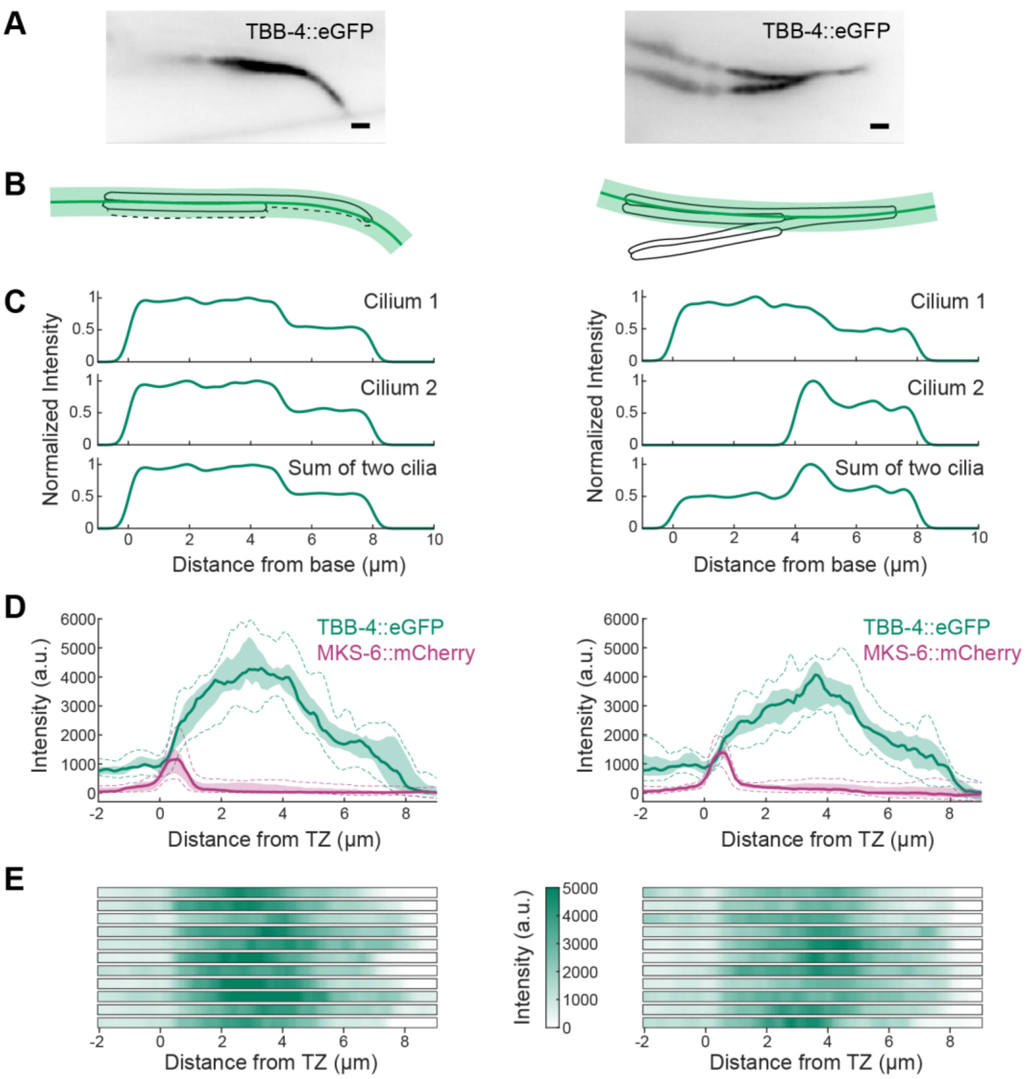


**Supplementary Figure 2: The influence of cilia pair orientation relative to the focal plane on the measured fluorescence intensity. (A)** Example inverted images of TBB-4::eGFP in the cilia, with two cilia spatially overlapping throughout the length (left) and for the last ~50% (right). Scale bar: 1 μm. **(B)** Schematic illustration of the contribution that cilium 2 adds to the intensity profile measured along cilium 1. Solid green line, spline drawn through the cilium; light green shading, line thickness chosen to cover the measured cilium completely. **(C)** Computer simulation of intensity profiles along cilium 1, cilium 2 and their sum (normalised to their maxima), in cases when cilia 1 and 2 overlap completely (left) and when the last 50% of cilium 2 overlaps with cilium 1. Note that the distance here is plotted starting from the ciliary base, which is located ~0.5 μm before the TZ. See Methods for the simulation details. **(D)** TBB-4::eGFP (green) intensity measured along one cilium, in the case of complete overlap of two cilia (left, n = 11 cilia pairs/worms) and when the proximal segments can be resolved (right, n = 11 cilia pairs/worms). MKS-6::mCherry (magenta) is used as a TZ marker. Solid lines, median; light shading, 25th-75th percentiles; dashed lines, 5th and 95th percentiles. **(E)** TBB-4::eGFP intensity distribution in individual cilia used in D.

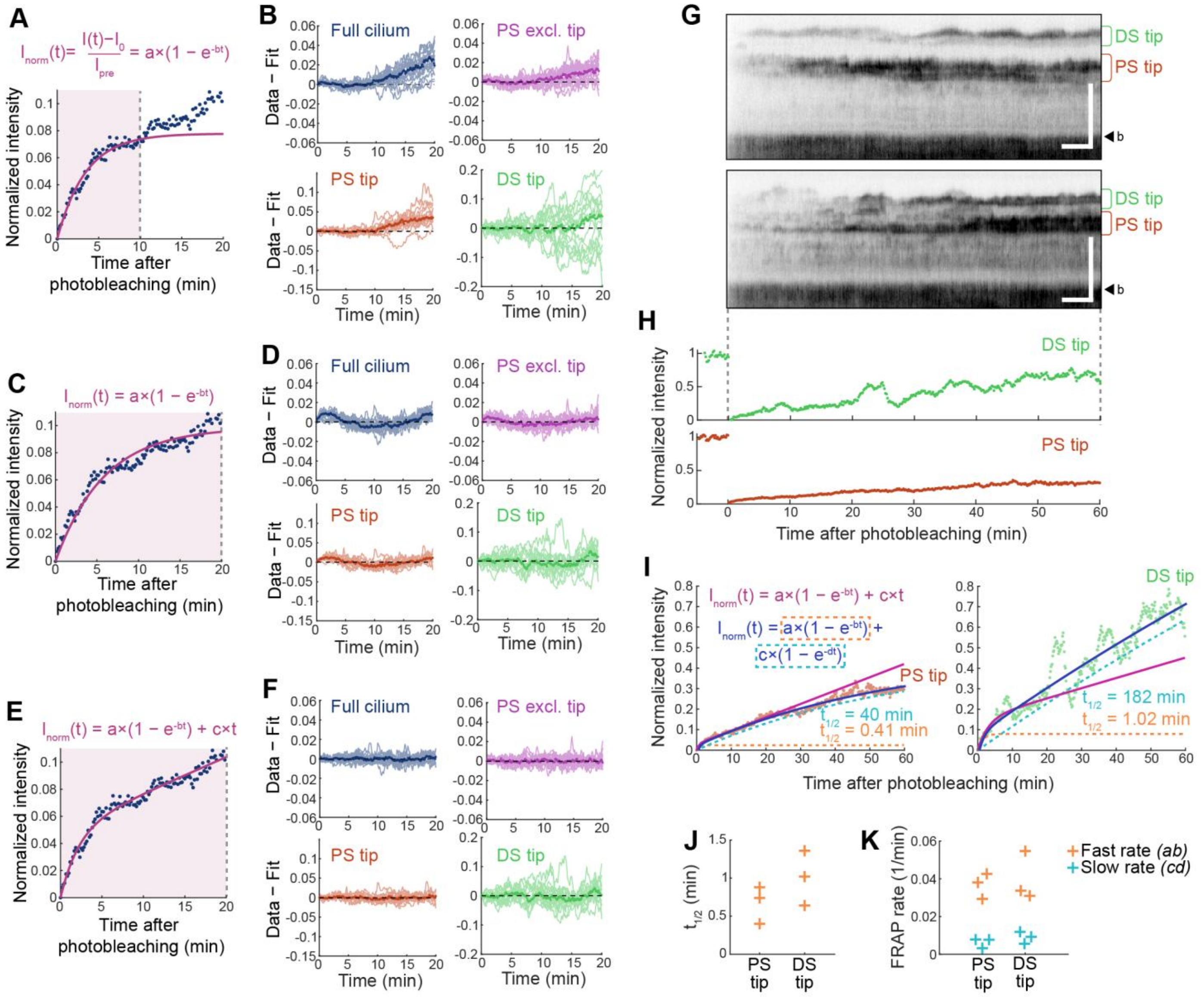


**Supplementary Figure 3: Fitting TBB-4 FRAP in the cilia. (A-F)** Evaluating different fitting options for TBB-4::eGFP FRAP curves. **(A, C, E)** Normalized TBB-4::eGFP intensity ($I_{norm}$) after photobleaching in the example full cilium shown in Figure 1D, H (blue dots) with first 10 minutes of fluorescence recovery fitted using a single-exponential equation **(A)**; 20 minutes of FRAP fitted using a single-exponential equation **(C)**; 20 minutes of FRAP fitted using an equation combining an exponential and a linear terms **(E)**. Fit result is plotted as a magenta line. Light shaded areas indicate which data points were used for the fit. **(B, D, F)** Residuals (Data - Fit) calculated using the fits shown in A, C, E, respectively, for the four regions in the cilia schematically illustrated in Figure 1G: Full cilium (blue), PS excluding the tip (magenta), PS tip (orange), DS tip (green). Dark lines represent the median, lighter lines – residuals from the individual worms (n = 17 for full cilium and PS excluding the tip; n = 15 for PS tip; n = 16 for DS tip). **(B)** Residuals increase at t>10 min, so the single exponential equation derived from fitting the first 10 minutes of FRAP cannot account for the full recovery process. **(D)** Residuals deviate from 0 in a similar manner for most worms, indicating that a single-exponential equation cannot correctly describe the observed FRAP. **(F)** The residuals stay close to 0 during the first 20 minutes of FRAP. On this time scale, the equation shown in E can reasonably well describe TBB-4::eGFP FRAP in the cilia. **(G)** Intensity-inverted kymographs for two examples of 1-hour-long FRAP. Scale bars: 5 µm (vertical), 5 min (horizontal); b, ciliary base. **(H)** Normalised intensity over time in the PS tip (orange) and DS tip regions (green) plotted for the bottom kymograph in G. Note the large intensity fluctuations at the DS tip. Those are also apparent from higher residual values for DS tip (light green lines in B, D, F). **(I)** Normalized intensity over time at PS tips (light orange dots) and DS tips (light green dots) shown in H fitted using a combination of exponential and linear terms based on first 20 minutes of FRAP (magenta line, similar to E) and with a combination of two exponential terms (blue line; dotted orange and light blue lines showing individual components). On an hour scale, linear term used in the first fit can no longer describe the FRAP. **(J)** Half-recovery time ($t_{1/2} = {ln2}/{b}$) of the fast exponent (orange dotted line in I) for PS and DS tip regions. While this value stays consistent between the worms (n = 3), half-recovery time of the second, slow exponent (light blue dotted line in I) varies widely (see $t_{1/2}$ examples in I) and likely cannot be reliably determined from our experiments. **(K)** Initial slopes of two exponents shown in I, measured in the PS and DS tips regions in n = 3 worms.

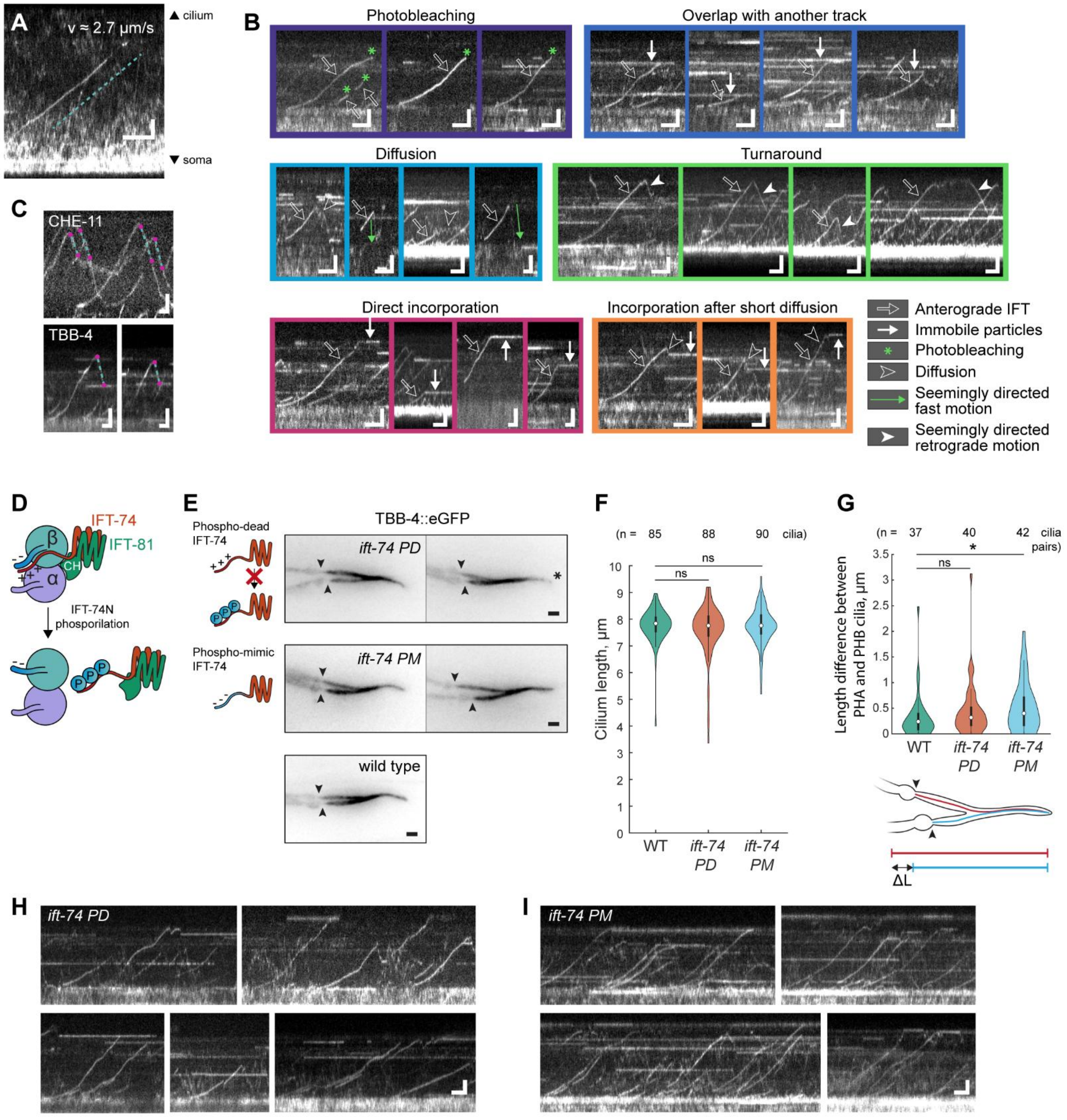


**Supplementary Figure 4: More examples of TBB-4 dynamics and their manual analysis. (A)** Kymograph showing a rare event of TBB-4 directed transport in the dendrite. The average slope of the track (dotted line) corresponds to velocity of 2.7 µm/s. Scale bars: 2 µm (vertical), 1 s (horizontal). **(B)** Examples of TBB-4 IFT tracks ending in different ways. Scale bars: 2 µm (vertical), 2 s (horizontal). **(C)** Illustration of retrograde motion velocity analysis. Two points (magenta dots) were selected on a linear-like region of retrograde directed tracks, and velocity was derived from the obtained slope. Scale bars: 2 µm (vertical), 2 s (horizontal). **(D)** A schematic depiction of mechanism of tubulin dissociation from IFT-B complex, suggested by Jiang, Shao et al. [1]. DYF-5-mediated phosphorylation of IFT-74 unstructured N-terminal domain decreases its net positive charge and weakens its ion interactions with negatively charged C-terminal tail of β-tubulin, which leads to dissociation of αβ-tubulin dimer from the IFT-B complex. **(E)** Inverted images (Z-stack maximum intensity projections) of TBB-4::eGFP in wild type background, phospho-dead (PD), and phospho-mimic (PM) *ift-74* mutants. In *ift-74 PD*, cilia mostly looked similar to wild type (left image), but in some cases the tip region appeared less clear, possibly indicating altered axoneme composition in this region (right image). In *ift-74 PM*, most cilia also looked similar to wild type (left image), but some PHA/PHB pairs had stronger misalignment of ciliary bases (right image). Arrowheads indicate ends of PCMC. Scale bar: 1 µm. **(F)** Violin plots of cilia lengths measured in wild type, *ift-74 PD* ($p = 0.30$), and *ift-74 PM* ($p = 0.42$), from the end of PCMC to the ciliary tip, using TBB-4::eGFP marker. To obtain a high number of cilia with all regions in focus, maximum-intensity projections of Z-stacks were used, so the determined length might be marginally lower than the actual length in all cases. On the other hand, since two cilia usually overlap in the DS, the tips of PHA and PHB cilia could not be detected independently and were determined by the longest of the two cilia. Statistical analysis was performed using nonparametric Mann-Whitney U test. **(G)** Violin plots of the difference between PHA and PHB cilia starting positions relative to the ciliary tip, measured using TBB-4::eGFP in wild type, *ift-74 PD* ($p = 0.067$), and *ift-74 PM* (*, $p = 0.026$). This difference was determined as the length difference between PHA and PHB cilia, measured from the end of their PCMC, to their common tip (see the schematic illustration below). Statistical analysis was performed using nonparametric Mann-Whitney U test. **(H, I)** Example kymographs showing TBB-4::eGFP dynamics in *ift-74 PD* (H) and *ift-74 PM* (I). Scale bars: 2 µm (vertical), 2 s (horizontal).

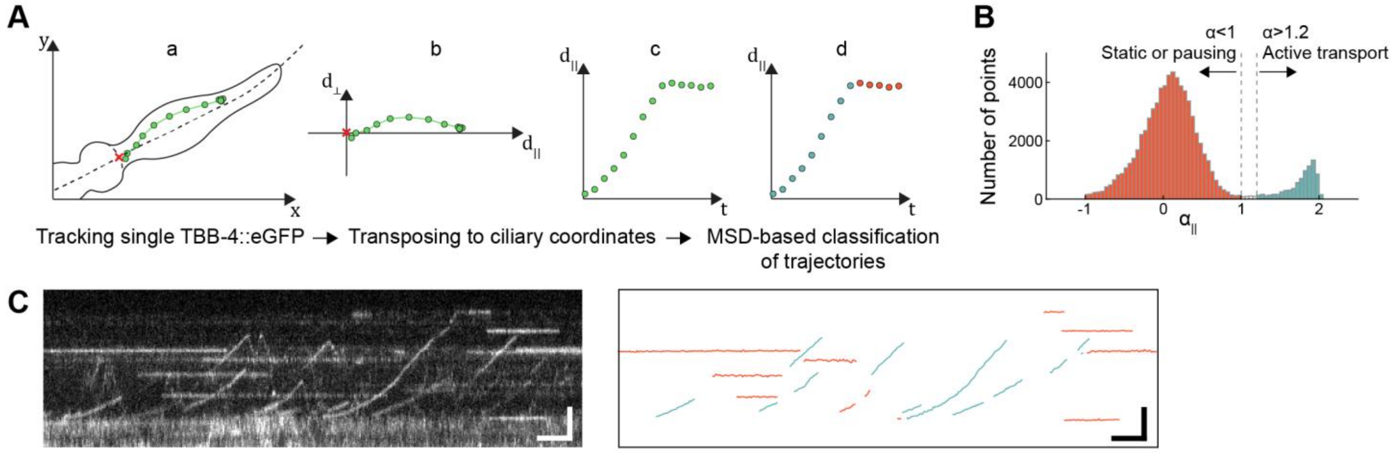


**Supplementary Figure 5: Automated trajectories classification and analysis. (A)** Analysis pipeline (also see Methods): (a) detecting localisations of single TBB-4::eGFP particles and connecting them into trajectories containing $x(t), y(t)$ information; (b) trajectories are transposed from Cartesian to cilium-based coordinates $d_{\parallel}(t)$, $d_{\perp}(t)$ by projecting them onto a hand-drawn spline through the middle of a cilium (dotted line in a); (c,d) using windowed mean squared displacement along the ciliary spline, (parts of) trajectories are classified as directed transport (d, blue) or static localisations (orange). **(B)** Distribution of $\alpha_{\parallel}$ derived from fitting the mean squared displacement along the spline for a window of 15 time frames for all trajectories as $\langle \Delta {d_{\parallel}}^2 \rangle (t) = \Gamma t^{\alpha}$. Since only IFT tracks and immobile particles were tracked at this frame rate, $\alpha_{\parallel}$ show a bimodal distribution. Trajectories fragments with α<1 were classified as static, α>1.2 as actively transported. **(C)** Example of trajectories detection and classification. Left: kymograph showing characteristic TBB-4::eGFP dynamics in the cilia. Right: trajectories extracted from this movie, classified as active transport (light blue) and static events (orange), as described in A-B. Scale bars: 2 µm (vertical), 2 s (horizontal).

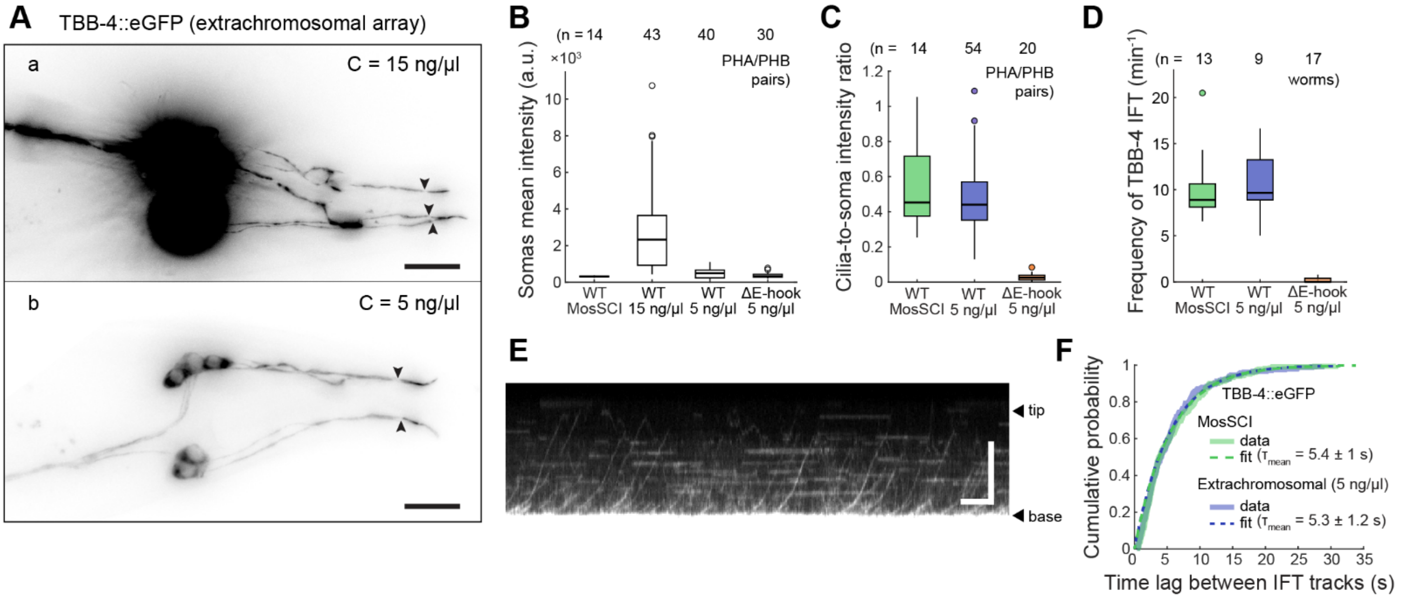


**Supplementary Figure 6: Observed differences in subcellular localisation and dynamics of ΔE-hook TBB-4::eGFP are not due to the differences in the expression levels. (A)** Inverted fluorescence microscopy images of tails of *C. elegans* expressing extrachromosomal tbb-4::egfp-coding construct (plasmid), microinjected at a concentration of 15 ng/µl (a) and 5 ng/µl (b). Higher construct concentration (a) results in strong overexpression and protein accumulation in the neurites. Lower plasmid concentration (b) results in an expression pattern similar to that in the MosSCI strain (Figure 1A). Scale bar: 10 µm. **(B)** TBB-4::eGFP expression level in the MosSCI and extrachromosomal strains, estimated as mean fluorescence intensity in the somas of PHA, PHB and PQR neurons. ΔE-hook TBB-4 strain used in this study (Figures 3, 4) has a similar level of labelled tubulin as a WT MosSCI strain. **(C)** Ratio between mean cilia and soma intensities in WT and ΔE-hook TBB-4::eGFP strains. The observed effect is caused not by differences in the expression level but by the ΔE-hook mutation. **(D)** Frequency of anterograde TBB-4 IFT tracks in WT and ΔE-hook TBB-4::eGFP strains. Tracks were manually picked from kymographs if they occurred in the first 1-2 µm of cilia (see Methods). **(E)** An example kymograph of ectopically expressed TBB-4::eGFP (C = 5 ng/µl). TBB-4 dynamics look similar to those in the MosSCI strain: active anterograde transport, diffusion, and incorporated particles. Scale bars: 5 µm (vertical), 5 s (horizontal). **(F)** Cumulative probability ($p$) distribution of time lags ($\tau$) between consecutive TBB-4 anterograde IFT tracks in MosSCI (n = 345 tracks) and extrachromosomal (n = 251 tracks) TBB-4::eGFP strains. Dotted lines, least square fit ($p = 1 - e^{-\lambda\tau}, \tau_{mean} = 1/\lambda$, mean and error estimated using bootstrapping).

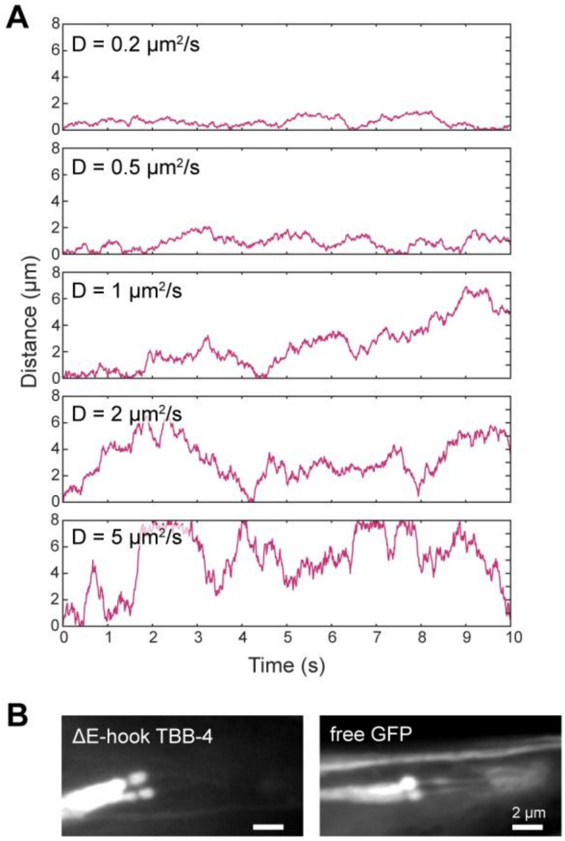


**Supplementary Figure 7: Interpreting TBB-4::eGFP diffusion. (A)** Examples of simulated diffusive tracks inside an 8 µm-long 'cilium'. When diffusion coefficients are high (a few $\mu m^2/s$), some parts of the tracks visually appear 'directed': particles can travel a distance of a few µm in one direction. However, it is still normal diffusion. **(B)** Fluorescence images of ΔE-hook TBB-4::eGFP (left) and eGFP (right) in PHA/PHB cilia. Both proteins are present in much higher amounts in the dendrites and PCMC than in the cilia.

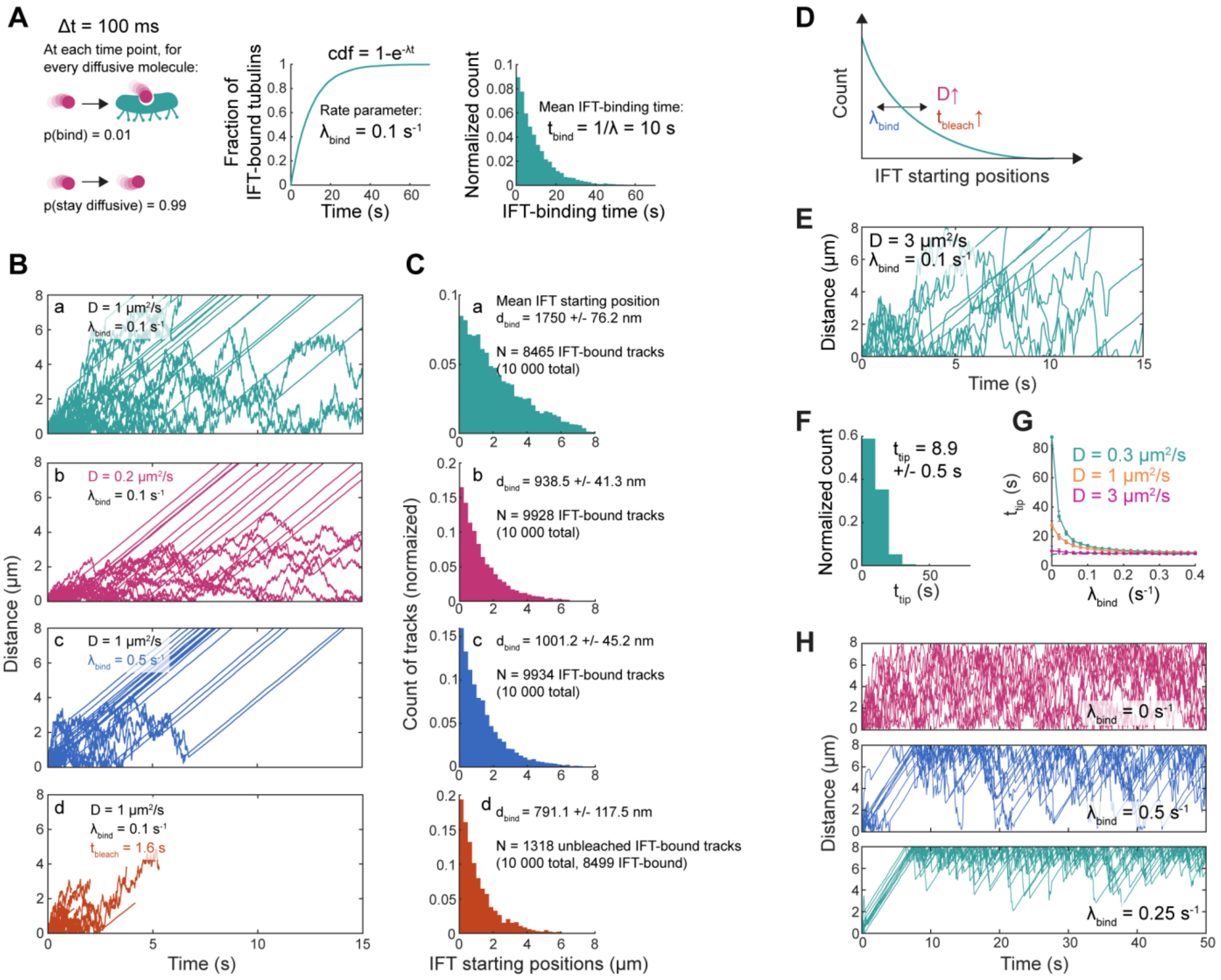


**Supplementary Figure 8: Assisting simulations to understand the role of IFT. (A)** Time step $\Delta t$ and probability $p$ to bind onto an IFT train at every time step are input parameters of the simulation. When many particles are analysed, the distribution of IFT-bound particles over time becomes exponential ($cdf = 1 - e^{-\lambda t}$, where λ is the rate parameter). The average time it takes for particles to bind to an IFT train is $t_{bind} = 1/\lambda$. **(B, C)** Related to Figure 2D. Simulated trajectories (B) and distributions of IFT-starting positions (C) obtained using different $D$, IFT-binding rate $\lambda_{bind}$ and photobleaching characteristic time $t_{bleach}$. When diffusive particles can bind IFT trains with equal probability at every time step, the distribution of IFT-binding positions is exponential (a). Increasing $\lambda_{bind}$ (c), decreasing $D$ (b) or $t_{bleach}$ (d) shift the distribution to the left. A combination of these factors can potentially explain the shape of the distribution we see in our data (Figure 2D). **(D)** A summary of factors influencing the distribution of IFT docking locations, as tested in C. **(E)** Example of simulated diffusive tracks ($D$ = 3 μm$^2$/s) when a small probability of binding to the IFT is present ($\lambda_{bind} = 0.1$ s$^{-1}$). Tracks stop being displayed once they reach the 'tip' ($x = 8$ μm). **(F)** Distribution of times it takes for particles to cover the distance from the base to the tip ($t_{tip}$) with $\lambda_{bind} = 0.1$ s$^{-1}$. **(G)** Dependency of $t_{tip}$ on IFT-binding rate $\lambda_{bind}$, obtained from simulations using different $D$ (n = 1000 tracks, mean and error calculated using bootstrapping). **(H)** Examples of simulated tracks ($D$ = 3 μm$^2$/s) with different IFT-binding rates. Here, tracks do not disappear after reaching the tip but get reflected back to the cilium. The distribution of particles in the cilia can be visually evaluated.

### Supplementary Tables

**Supplementary Table 1. C. elegans strains used in this study**

| Strain | Genotype | Source | Short notation |
|---|---|---|---|
| EJP401 | *vuaSi401 [pSA401; Ptbb- 4::tbb-4::eGFP; cb-unc- 119(+)] I* | Oswald et al., 2018 [2] | TBB-4 |
| EJP601 | *vuaSi401 [pSA401; Ptbb- 4::tbb-4::eGFP; cb-unc- 119(+)] I; vuaSi21 [pBP39; Pmks-6::mks-6::mCherry; cb-unc-119(+)] II* | This study | TBB-4, MKS-6 |
| EJP602 | *vuaSi401 [pSA401; Ptbb- 4::tbb-4::eGFP; cb-unc- 119(+)] I; vuaSi24 [pBP43; Pche-11::che-11::mCherry; cb-unc-119(+)] II; che-11(tm3433) V* | This study | TBB-4, CHE-11 |
| EJP603 | Ex[*Ptbb- 4::tbb-4::eGFP; cb-unc- 119(+), myo-2::mCherry*] | This study | TBB-4 (extrachromosomal array) |
| EJP604 | Ex[*Ptbb- 4::tbb-4(ΔE-hook)::eGFP; cb-unc-119(+), myo-2::mCherry*] | This study | ΔE-hook TBB-4 (extrachromosomal array) |
| EJP605 | *vuaSi24 [pBP43; Pche-11::che-11::mCherry; cb-unc-119(+)] II; che-11(tm3433) V;* Ex[*Ptbb-4::tbb-4(ΔE-hook)::eGFP; cb-unc- 119(+), myo-2::mCherry*] | This study | ΔE-hook TBB-4 (extrachromosomal array), CHE-11 |
| AML10 | otls355 [rab-3::NLS::tagRFP]; otls45 [unc-119::GFP] V | Nguyen et al., 2016 [3] | Free GFP |
| PHX7406 | vuaSi401 I; ift-74 (syb4674) [ift-74 PM knock-in] II | This study (SunyBiotech) | TBB-4, ift-74 PM |
| PHX7429 | vuaSi401 I; ift-74 (syb4679) [ift-74 PD knock-in] II | This study (SunyBiotech) | TBB-4, ift-74 PD |

**Supplementary Table 2. Fast and slow components of TBB-4 FRAP** (mean $\pm$ 3σ obtained from bootstrapping)

| | Full cilium | PS excl. the tip | PS tip | DS tip |
|---|---|---|---|---|
| Fast initial rate ($a \cdot b$, $min^{-1}$) | $(2.89 \pm 0.70) \cdot 10^{-2}$ | $(2.49 \pm 0.48) \cdot 10^{-2}$ | $(3.76 \pm 1.18) \cdot 10^{-2}$ | $(3.58 \pm 2.32) \cdot 10^{-2}$ |
| Fast $t_{1/2}$ ($ln2/b$, min) | $1.07 \pm 0.34$ | $1.42 \pm 0.38$ | $0.87 \pm 0.35$ | $2.68 \pm 9.89$ |
| Slow rate ($c$, $min^{-1}$) | $(2.3 \pm 0.85) \cdot 10^{-3}$ | $(1.2 \pm 0.72) \cdot 10^{-3}$ | $(3.1 \pm 1.9) \cdot 10^{-3}$ | $(4.7 \pm 5.0) \cdot 10^{-3}$ |
| N, worms | 17 | 17 | 15 | 16 |

**Supplementary Table 3. Sequence and primers used for ΔE-hook TBB-4::eGFP construct**

| Sequence, 5'-3' | Explanation |
|---|---|
| **QQYQEATADDEGEFDEHDQDVE**AGGGSGGGGSGGGGS | Amino acid sequence of TBB-4 E-hook (bold), followed by a NgoMIV restriction site and Gly-Ser linker connecting TBB-4 and eGFP in our TBB-4::eGFP construct. The highlighted region was removed in the ΔE-hook strain. |
| CGTAATACGACTCACTAGTGGGCAGATCTACTATTTCTGTAACTACGCG | Forward primer for Ptbb-4::tbb-4(ΔE-hook) amplification with an overlap with the pCFJ350 region upstream of the BglII restriction site, for Gibson assembly |
| CACCTCCTCCGCTTCCTCCGCCGGCGGCGGTTGCTTCTTGATACTGTTG | Reverse primer for Ptbb-4::tbb-4(ΔE-hook) amplification with an overlap with NgoMIV site and linker, for Gibson assembly |
| CGCCCAGGAGAACACGTTAG | Forward sequencing primer to confirm Ptbb-4::tbb-4(ΔE-hook) correct ligation |

| | |
|---|---|
| CCGTATGTTGCATCACCTTCAC | Reverse sequencing primer to confirm Ptbb-4::tbb-4(ΔE-hook) correct ligation |

### Supplementary Notes

**Supplementary Note 1. Fitting TBB-4::eGFP FRAP curves**

We first fitted our TBB-4::eGFP FRAP time trace to a single exponential function, similar to an earlier FRAP study where fluorescence recovery was monitored over 10 min [4]. It yielded satisfactory fits only when the first 10 minutes of the data were considered (Supplementary Figure 3A-B), but failed for the complete 20-minute traces (Supplementary Figure 3C-D). This indicates that, apart from a component decaying on the minute time scale, the data also contains an additional component decaying far more slowly. Double-exponential fits did not work since the measurement time window was too short to reliably fit the additional slow recovery component. To overcome this problem, we tried a combination of an exponential and a linear term: $I_{norm}(t) = \frac{I(t)-I_0}{I_{pre}} = a\left(1 - e^{-bt}\right) + ct$, where $I(t)$ is the time-dependent mean background-subtracted fluorescence intensity in the ROI, $I_0$ is fluorescence intensity right after photobleaching, $I_{pre}$ – before photobleaching, (Figure 1I, Supplementary Figure 3E-F). The exponential term (Figure 1I, orange line; characterized by parameters $a$ and $b$, with slope at $t = 0$, equal to $a \cdot b$) dominates during the first few minutes ($t_{1/2} \sim$ 1-2 min, see Supplementary Table 2), while the linear term dominates the long-time behaviour (Figure 1I, light blue line; characterised by slope $c$).

Our fitting results suggest that fluorescence recovery happens at two different rates ($a \cdot b$ and $c$), differing by an order of magnitude. In the initial exponential phase, PS tips recover fastest, followed by the DS tips, possibly reflecting the differences in proximity of these regions to the dendrite. DS tips show the highest rate in the later linear-like phase, followed by the PS. This could reflect the range in dynamic instability of A- versus B-tubules: the further MTs shrink during catastrophe events, the more fluorescent tubulin can get incorporated during the slow recovery phase. In both cases, the DS tip region has the largest error arising from the large intensity fluctuations (Figure 1H). We note that the fit function we used cannot describe the full recovery process, since the fluorescence intensity cannot increase infinitely with time (as predicted by the fit function). This is the simplest approximation that can be exploited to describe the first 20 minutes after photobleaching. On longer time scales, the linear term $ct$ , most likely turns into an exponential term. Evidence for this was obtained by double exponential fits of the few successful 1-hour recovery curves, which appeared to reach an equilibrium (Supplementary Figure 3G-I). We note, however, that we could only obtain a few of such movies, since worms rarely stayed still and in focus throughout the experiment. In conclusion, FRAP data of TBB-4::eGFP in phasmid cilia show that the recovery dynamics of microtubules within cilia are complex, taking place on characteristic time scales of about a minute and tens of minutes.

**Supplementary Note 2. Reasoning behind the choice of the TBB-4 strains**

We used a MosSCI WT TBB-4 strain with the original copy of the tbb-4 gene unaffected to avoid excessive concentrations of fluorescent protein that could interfere with tubulin function. For the ΔE-hook mutant, we used an extrachromosomal array, mainly for practical convenience reasons. To be able to quantitatively compare the WT and ΔE-hook strains, we compared TBB-4::eGFP expression in their phasmid neurons and chose the strains in which expression levels were similar (Supplementary Figure 6B). To check if the way of delivering the construct did not affect the protein function, we also created a WT TBB-4::eGFP extrachromosomal array, with an expression level similar to that of the MosSCI strain, and imaged TBB-4 dynamics in it (Supplementary Figure 6E). The frequency of TBB-4 IFT was similar in both MosSCI and extrachromosomal array strains, but was significantly lower in the ΔE-hook TBB-4 strain (Supplementary Figure 6D, F).

**Supplementary Note 3. Interpreting directed diffusive tracks**

We wondered whether the fast, apparently directional motion that we sometimes observed in cilia and dendrites (Figure 2B, 3F) can be explained by normal diffusion or requires some other mechanisms. We simulated particles diffusing with different diffusion coefficients in one dimension along an 8 µm-long 'cilium' (Supplementary Figure 7A). We noted that at $D$>2 µm$^2$/s, occasionally, tracks in the kymographs appear directed for several seconds. It could well be that our analysis is positively biased in favour of such 'directed' tracks: when many particles simultaneously diffuse into the field of view, the rare trajectory with multiple consecutive 'steps' in one direction would escape the busy region with many overlapping tracks, and therefore be the only one that can be tracked using our algorithms. In conclusion, it is not unexpected that a fraction of the trajectories of fast diffusing particles appear 'directed'. Such directed tracks cannot be considered proof for additional fast transport mechanisms, different from IFT, but do not exclude this possibility. Similar tracks have been observed before for α-tubulin [5] and EB1 [6] in *C. reinhardtii*.

### Supplementary Movies

**Supplementary Movie 1.** Three stages of the TBB-4::eGFP FRAP experiment. Left panel, time-lapse imaging before photobleaching (5 min). Middle panel, photobleaching (5 min). Right panel, time-lapse imaging after photobleaching (20 min). Brightness settings for the left and right panels are the same. Imaging conditions are: for time-lapse imaging – 1 frame every 10 s, 150 ms exposure time, 491 nm excitation at ~0.2 W/mm$^2$, for photobleaching – continuous imaging at 50 fps, 491 nm excitation at ~10 W/mm$^2$.

**Supplementary Movie 2.** Examples of TBB-4:eGFP FRAP in different worms. Left panel, time-averaged projection before photobleaching. Right panel, fluorescence recovery (1 hour or 20 min-long). Brightness settings for the left and right panels are the same.

**Supplementary Movie 3.** Examples of TBB-4::eGFP dynamics in the dendrites of phasmid neurons in 4 different worms. Left panel, fragments of movies acquired at 37-46 fps. Right panel, time-average projections.

**Supplementary Movie 4.** Examples of TBB-4::eGFP dynamics in phasmid cilia in 2 different worms. Left panel, fragments of movies acquired at 13 and 19.3 fps, respectively. Right panel, time-average projections.

**Supplementary Movie 5.** Simultaneous imaging of CHE-11 (IFT-A) and TBB-4 in phasmid cilia. Top panel, TBB-4::eGFP. Middle panel, CHE-11:mCherry. Bottom panel, merged view. The movie is acquired with an exposure time of 51.9 ms per frame, using alternating illumination with 491 and 561 nm lasers, resulting in a 103.8 ms time interval per frame for each channel. The video is played at 2x speed.

**Supplementary Movie 6.** Examples of ΔE-hook TBB-4::eGFP dynamics in the phasmid cilia of 3 different worms. Left panel, fragments of movies acquired at 19.3 fps. Right panel, maximum intensity projections.
